# Analysis of FGF20-regulated genes in organ of Corti progenitors by translating ribosome affinity purification

**DOI:** 10.1101/2020.04.13.040212

**Authors:** Lu M. Yang, Lisa Stout, Michael Rauchman, David M. Ornitz

## Abstract

**Background:** Understanding the mechanisms that regulate hair cell (HC) differentiation in the organ of Corti (OC) is essential to designing genetic therapies for hearing loss due to HC loss or damage. We have previously identified Fibroblast Growth Factor 20 (FGF20) as having a key role in HC and supporting cell differentiation in the mouse OC. To investigate the genetic landscape regulated by FGF20 signaling in OC progenitors, we employ Translating Ribosome Affinity Purification combined with Next Generation mRNA Sequencing (TRAPseq) in the *Fgf20* lineage.

**Results:** We show that TRAPseq targeting OC progenitors effectively enriched for mRNA within this rare cell population. TRAPseq identified differentially expressed genes downstream of FGF20, including *Etv4, Etv5, Etv1, Dusp6, Hey1, Hey2, Heyl, Tectb, Fat3, Cpxm2, Sall1, Sall3*, and cell cycle regulators such as *Cdc20*. Analysis of *Cdc20* conditional-null mice identified decreased cochlea length, while analysis of *Sall1-ΔZn*^*2-10*^ mice, which harbor a mutation that causes Townes-Brocks syndrome, identified a decrease in outer hair cell number.

**Conclusions:** We present two datasets: genes with enriched expression in OC progenitors, and genes regulated by FGF20 in the embryonic day 14.5 cochlea. We validate select differentially expressed genes via in situ hybridization and in vivo functional studies in mice.

**Key findings:**

- Translating Ribosome Affinity Purification (TRAP) with Fgf20-Cre enriches for prosensory cell mRNA
- TRAP combined with RNAseq identifies genes downstream of FGF20 during prosensory cell differentiation
- FGF20 regulates Sall1, gene implicated in human sensorineural hearing loss

**Grant Sponsor and Number:** National Institute on Deafness and Other Communication Disorders – DC017042 (DMO) Washington University Institute of Clinical and Translational Sciences and National Center for Advancing Translational Sciences – CTSA grant UL1TR002345 (JIT471 to DMO) March of Dimes – 6-FY13-127 (MR)

## INTRODUCTION

Congenital and acquired sensorineural hearing loss are common problems, yet there are no available biologically-based therapies. Congenital sensorineural hearing loss can result from defects in sensory hair cells (HCs) or specialized supporting cells (SCs) within the organ of Corti (OC) (Allen and Goldman 2019; Basch et al. 2016; Bowl and Brown 2018; Wu and Kelley 2012). Acquired sensorineural hearing loss is commonly caused by damage to HCs (Wong and Ryan 2015; Yamasoba et al. 2013). In mammals, HC loss is permanent as the mammalian OC is unable to regenerate HCs (Corwin and Warchol 1991; Wong and Ryan 2015). One potential approach to treating hearing loss due to HC loss or damage is to reactivate developmental signaling pathways in latent progenitors to promote their growth and differentiation into HCs and SCs. Investigation of the developmental pathways regulating HC and SC differentiation will benefit our understanding and treatment of both congenital and acquired hearing loss.

In mouse cochlea development, Fibroblast Growth Factor 20 (FGF20) signaling via FGF receptor 1 (FGFR1) is required for the differentiation of organ of Corti progenitors (prosensory cells) into HCs and SCs, specifically outer hair cells (OHCs) and outer supporting cells (Hayashi et al. 2008; Huh et al. 2012, 2015; Ono et al. 2014; Pirvola et al. 2002). *Fgf20*-null mice (*Fgf20*^*-/-*^) are deaf, with loss of OHCs and gaps of undifferentiated cells along the length of the OC interrupting the normal patterning of HCs and SCs (Huh et al. 2012). *Fgf20*^*-/-*^ cochleae also exhibit shorter cochlear length. Additionally, FGF20 is required during the initiation of HC and SC differentiation and *Fgf20*^*-/-*^ mice have premature onset of HC differentiation, as well as delayed apical progression of HC differentiation and maturation (Huh et al. 2012; Yang et al. 2019). However, we do not know the mechanism by which FGF20 is required for the initiation of differentiation. We hypothesize that downstream genetic targets of FGF20 signaling in prosensory cells will be candidate effectors of HC and SC differentiation. Identifying these genes will be important for advancing therapeutics in regenerating lost or damaged HCs and will provide insight into the mechanisms underlying OC phenotypes in *Fgf20*^*-/-*^ mice.

Here, we combined the Translating Ribosome Affinity Purification (TRAP) technology (Heiman et al. 2008) with Next Generation mRNA Sequencing (TRAPseq) to study changes in gene expression patterns in prosensory cells in the presence or absence of FGF20 signaling. TRAP allows the isolation of translating mRNA from specific cell populations without cell sorting or fine dissection. We used the *ROSA*^*fsTRAP*^ allele (Zhou et al. 2013), which when activated by Cre recombinase, leads to the expression of a GFP-tagged ribosomal protein (L10a-eGFP). Immunoprecipitation (IP) for GFP then isolates polysomes and associated translating mRNA. We show that by targeting the expression of L10a-eGFP to prosensory cells within the cochleae, we were able to enrich for translating mRNA within this relatively rare cell population. Comparing control and *Fgf20*^*-/-*^ prosensory cell mRNA, TRAPseq revealed many genes previously associated with FGF signaling, as well as genes with functional significance in cochlea development. Among these genes is *Sall1*, mutations in which cause Townes-Brocks syndrome, a genetic condition associated with variable features that include sensorineural hearing loss (Kohlhase et al. 1998; Rossmiller and Pasic 1994).

## RESULTS

### *Fgf20*^*Cre*^ targets L10a-eGFP expression to the cochlear prosensory domain and Kölliker’s organ

At embryonic day 14.5 (E14.5), the floor of the cochlear duct can be divided into three sections (Fig. 1A): 1) prosensory domain (PD), which contains prosensory cells that differentiate into HCs and SCs of the OC; 2) outer sulcus (OS), epithelium that is lateral (abneural) to the prosensory domain, which develops into the lesser epithelial ridge (LER), and 3) Kölliker’s organ (KO), epithelium that is medial (neural) to the prosensory domain, which develops into the greater epithelial ridge (GER). We have previously shown that at E14.5, *Fgf20* is expressed in the prosensory domain and at postnatal day 1 (P1), the *Fgf20*^*Cre*^ lineage includes the OC and the GER (Huh et al. 2012, 2015).

**Figure 1.**
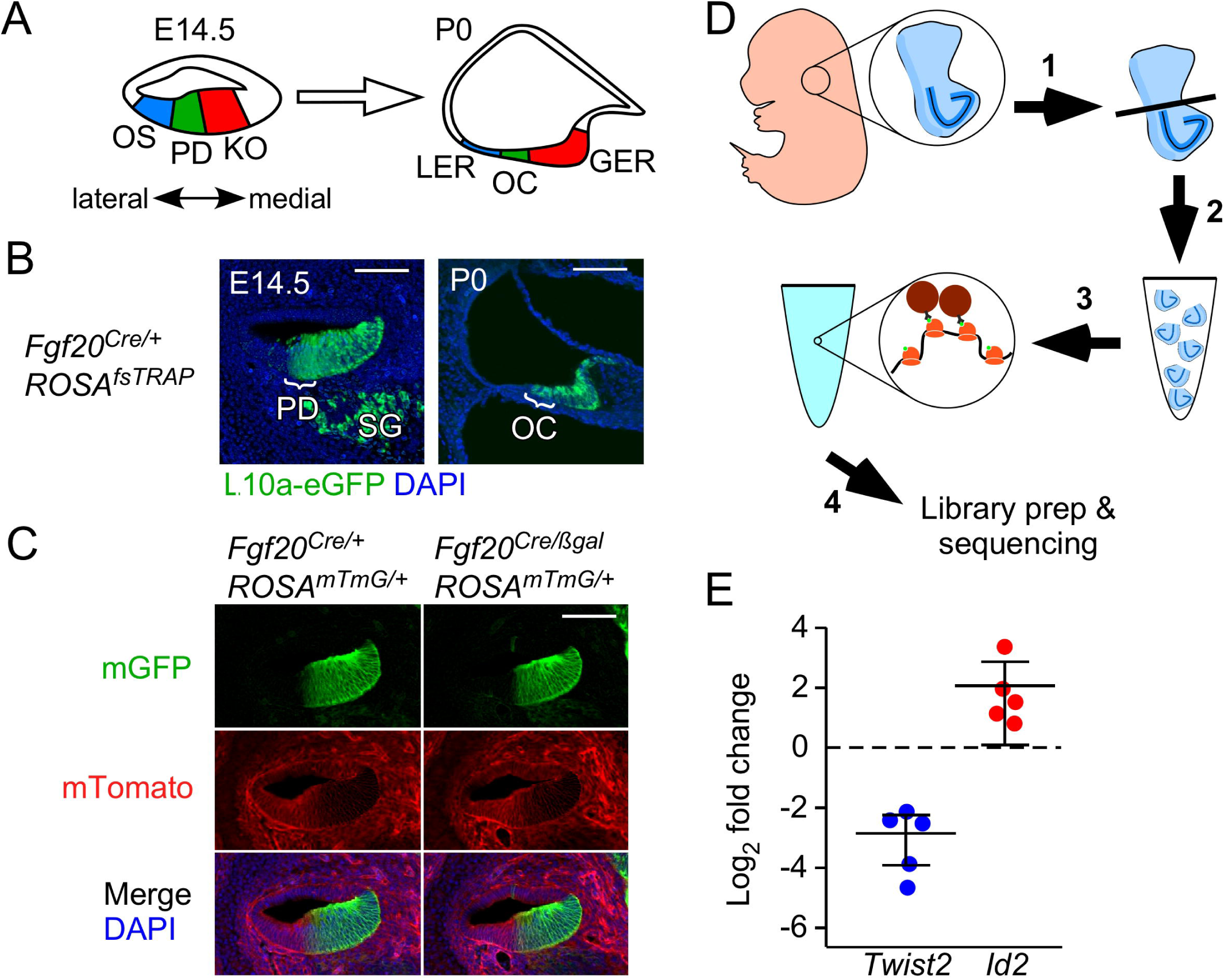
*Fgf20*^*Cre*^ targets L10a-eGFP expression to the cochlear prosensory domain and Kölliker’s organ. (A) Schematic representing cross-sectional view through the E14.5 and P0 cochlear duct. At E14.5, the epithelium at the cochlear duct floor can be divided into three regions: outer sulcus (OS), prosensory domain (PD), and Kölliker’s organ (KO). Cells from these three regions contribute to the lesser epithelial ridge (LER), organ of Corti (OC), and greater epithelial ridge (GER), respectively, at P0. Double-headed arrow indicates medial (neural) and lateral (abneural) directions. (B) Sections through the middle turn of E14.5 and P0 *Fgf20*^*Cre/+*^;*ROSA*^*fsTRAP/+*^ cochlear ducts, showing L10a-eGFP (green) expression. At E14.5, L10a-eGFP is found in the prosensory domain (PD; bracket), Kölliker’s organ and medial wall, and spiral ganglion (SG). At P0, it is found in the organ of Corti (OC; bracket) and greater epithelial ridge. DAPI, nuclei (blue); scale bar, 100 µm. (C) Section through the middle turn of E14.5 cochlear ducts from *Fgf20*^*Cre/+*^;*ROSA*^*mTmG/+*^ and *Fgf20*^*Cre/βgal*^;*ROSA*^*mTmG/+*^ embryos. Cells of the *Fgf20*^*Cre*^-lineage express mGFP (mG, green); non-lineage cells express mTomato (mT, red). DAPI, nuclei (blue); scale bar, 12 100 µm. (D) Schematic showing an overview of the TRAPseq protocol (see Experimental Procedures). 1) Ventral otocysts containing the cochlea were dissected from E14.5 embryos. 2) Otocysts from each litter were pooled according to genotype to increase RNA yield. 3) Otocysts were then homogenized and centrifuged to make polysomes before immunoprecipitation with anti-GFP antibodies to collect L10a-eGFP labelled polysomes, 4) which were then used for downstream applications. (E) qRT-PCR showing fold change in *Twist2* and *Id2* expression (normalized to *Gadph*) in TRAP RNA samples compared to pre-TRAP samples from *Fgf20*^*Cre/+*^;*ROSA*^*mTmG/+*^ E14.5 cochleae pooled from at least three embryos. Each dot represents a pooled sample.

To evaluate the TRAP technique for our use, we combined the *ROSA*^*fsTRAP*^ and *Fgf20*^*Cre*^ alleles. The *Fgf20*^*Cre*^ allele was made by targeted insertion of a sequence encoding a GFP-Cre fusion protein replacing exon 1 of *Fgf20* (Huh et al. 2015). As expected based on prior expression and lineage tracing experiments, at E14.5, L10a-eGFP fluorescence (green) from *Fgf20*^*Cre/+*^; *ROSA*^*fsTRAP*^ cochleae was found in the prosensory domain, Kölliker’s organ, the cochlear duct wall more medial to the Kölliker’s organ, and some cells in the spiral ganglion (Fig. 1B). Also, as expected, at P0, L10a-eGFP in *Fgf20*^*Cre/+*^; *ROSA*^*fsTRAP*^ cochleae was found in the OC and the GER (Fig. 1B).

Another *Fgf20* null allele, *Fgf20*^*βgal*^, was made by targeted insertion of a sequence coding β-Galactosidase replacing exon 1 of *Fgf20* (Huh et al. 2012). We combined the *Fgf20*^*Cre*^ and *Fgf20*^*βgal*^ alleles to generate *Fgf20*^*-/-*^ mice (*Fgf20*^*Cre/βgal*^), which maintained the same dosage of Cre as control mice (*Fgf20*^*Cre/+*^). Importantly, based on both double fluorescence expression from the *ROSA*^*mTmG*^ Cre-reporter allele, the *Fgf20*^*Cre*^ lineage (green) did not change in *Fgf20*^*Cre/βgal*^ cochleae compared to *Fgf20*^*Cre/+*^ (Fig. 1C). Based on these results, we believed that *Fgf20*^*Cre/+*^; *ROSA*^*fsTRAP/+*^ and *Fgf20*^*Cre/βgal*^; *ROSA*^*fsTRAP/+*^ mice will allow us to enrich for prosensory cell mRNA to examine changes in gene expression in the absence of FGF20 signaling.

### *Fgf20*^*Cre*^ TRAPseq enriched for prosensory domain mRNA

TRAPseq experiments were performed at E14.5, based on our previous findings that FGF20 signaling is required for prosensory cell differentiation at E13.5-E15.5 (Yang et al. 2019). In the initial experiment, we collected pre-TRAP (pre-IP) and TRAP (post-IP) RNA from *Fgf20*^*Cre/+*^;*ROSA*^*fsTRAP/+*^ cochleae at E14.5 (Fig. 1D). Quantitative reverse transcription PCR (qRT-PCR) showed enrichment of the prosensory cell marker *Id2* (Jones et al. 2006) and depletion of the mesenchyme marker *Twist2* (also called *Dermo1*) (Huh et al. 2015), by TRAP 5 (Fig. 1E).

Next, we performed TRAPseq on *Fgf20*^*-/+*^ (*Fgf20*^*Cre/+*^; *ROSA*^*fsTRAP/+*^) control and *Fgf20*^*-/-*^ (*Fgf20*^*Cre/βgal*^; *ROSA*^*fsTRAP/+*^) E14.5 cochleae. *Fgf20*^*-/+*^ and *Fgf20*^*-/-*^ embryos were generated at a 1:1 ratio. For each litter, cochleae from all control embryos were pooled together for RNA collection, and likewise for *Fgf20*^*-/-*^ embryos. Each sample represents RNA from pooled tissue from a minimum of three embryos. In total, 24 libraries were sequenced: 16 TRAP samples (8 *Fgf20*^*-/+*^ and 8 *Fgf20*^*-/-*^) and 8 pre-TRAP samples (4 *Fgf20*^*-/+*^ and 4 *Fgf20*^*-/-*^). Pre-TRAP samples were collected prior to IP, representing whole cochlea RNA, including RNA from mesenchyme and otic capsule. See Experimental Procedures for details.

Principal component analysis (PCA) of the 24 samples showed separation between pre-TRAP and TRAP RNA samples along PC1 (Fig. 2A). However, there was no separation between *Fgf20*^*-/+*^ vs. *Fgf20*^*-/-*^ samples along PC1 or PC2. PCA of only the 16 TRAP samples also did not show separation between *Fgf20*^*-/+*^ vs. *Fgf20*^*-/-*^ samples along the first two PCs (Fig. 2B). To assess the efficiency of the TRAP technique, differentially expressed gene (DEG) analysis using DESeq2 (Love et al. 2014) was performed to compare pre-TRAP control samples with TRAP control samples (same genotype, *Fgf20*^*Cre/+*^;*ROSA*^*fsTRAP/+*^, for both). 3850 DEGs were identified with adjusted p-value (padj) < 0.01 and Log_2_ Fold Change (LFC) < −1 or > 1 (Fig. 2C). Of these, 2017 genes had decreased expression in TRAP samples, compared to pre-TRAP (depleted by TRAP) and 1833 genes had increased expression in TRAP samples, compared to pre-TRAP (enriched by TRAP). Among the genes depleted by TRAP were mesenchymal markers *Cd44* and *Twist2* (Huh et al. 2015; Zhu et al. 2006), vasculature markers *Eln* and *Fbln1* (Cooley et al. 2008; Karnik et al. 2003), and chondrocyte markers *Runx2* and *Matn1* (Fujita et al. 2004; Pei et al. 2008). This was expected, since otic mesenchyme and capsule were included in the input tissue but did not express L10a-eGFP. *Bmp4, Lmx1a*, and *Gata2*, markers for the outer sulcus (Lilleväli et al., 2004; Ohyama et al., 2010) were depleted as well. This was also expected, as the outer sulcus was not captured by TRAP (Fig. 1A, B). Among the genes enriched by TRAP were prosensory domain markers *Fgf20, Atoh1, Hey2, Sox2, Gata3*, and *Id2* (Basch et al. 2011; Huh et al. 2012; Jones et al. 2006; Kiernan et al. 2005, 9; Luo et al. 2013; Woods et al. 2004), Kölliker’s organ markers *Lfng, Fgf10*, and *Jag1* (Ohyama et al. 2010), and spiral ganglion markers *Neurod1* and *Tubb3* (Locher et al. 2014; Puligilla et al. 2010), as expected based on the *Fgf20*^*Cre*^ lineage. Gene set overlap analysis with gene ontology (GO) on genes depleted by TRAP showed biological processes terms “angiogenesis” and “endochondral ossification” among the top terms (Table 1). GO analysis on genes enriched by TRAP showed biological processes terms “sensory perception of sound”, “axon guidance”, and “auditory receptor cell stereocilium organization” among the top terms (Table 2). These results strongly suggest that TRAP enriched for RNA from our target cell population. Pre-TRAP vs. TRAP sequencing comparison data are presented in Supplemental file S1.

**Table 1.** Top 12 enriched gene ontology (GO) terms from a list of 2017 differentially expressed genes depleted by TRAP, compared to pre-TRAP samples.

| Rank* | GO ID | GO biological processes term | p-value |
| --- | --- | --- | --- |
| 1 | GO:0006954 | inflammatory response | 3.30E-15 |
| 2 | GO:0001525 | angiogenesis | 1.60E-12 |
| 3 | GO:0030198 | extracellular matrix organization | 2.20E-11 |
| 4 | GO:0045766 | positive regulation of angiogenesis | 2.20E-10 |
| 5 | GO:0070374 | positive regulation of ERK1 and ERK2 cascade | 2.70E-10 |
| 6 | GO:0001974 | blood vessel remodeling | 3.20E-10 |
| 7 | GO:0007155 | cell adhesion | 8.80E-10 |
| 8 | GO:0030593 | neutrophil chemotaxis | 2.00E-09 |
| 9 | GO:0002548 | monocyte chemotaxis | 8.70E-09 |
| 10 | GO:0007186 | G-protein coupled receptor signaling pathway | 5.10E-08 |
| 11 | GO:0090090 | negative regulation of canonical Wnt signaling pathway | 2.20E-07 |
| 12 | GO:0001958 | endochondral ossification | 2.80E-07 |

\* Rank by p-value (lowest to highest).

**Table 2.** Top 12 enriched gene ontology (GO) terms from a list of 1833 differentially expressed genes enriched by TRAP, compared to pre-TRAP samples.

| **Rank\*** | **GO ID** | **GO biological processes term** | **p-value** |
| --- | --- | --- | --- |
| 1 | GO:0007605 | sensory perception of sound | 6.30E-09 |
| 2 | GO:0007411 | axon guidance | 3.30E-08 |
| 3 | GO:0048791 | calcium ion-regulated exocytosis of neurotransmitter | 7.00E-08 |
| 4 | GO:0048172 | regulation of short-term neuronal synaptic plasticity | 1.20E-06 |
| 5 | GO:0042391 | regulation of membrane potential | 2.00E-06 |
| 6 | GO:0007626 | locomotory behavior | 3.00E-06 |
| 7 | GO:0050885 | neuromuscular process controlling balance | 3.60E-06 |
| 8 | GO:0019228 | neuronal action potential | 7.20E-06 |
| 9 | GO:0014059 | regulation of dopamine secretion | 9.90E-06 |
| 10 | GO:0017158 | regulation of calcium ion-dependent exocytosis | 1.10E-05 |
| 11 | GO:0060088 | auditory receptor cell stereocilium organization | 1.70E-05 |
| 12 | GO:0045665 | negative regulation of neuron differentiation | 2.30E-05 |

\* Rank by p-value (lowest to highest).

**Figure 2.**
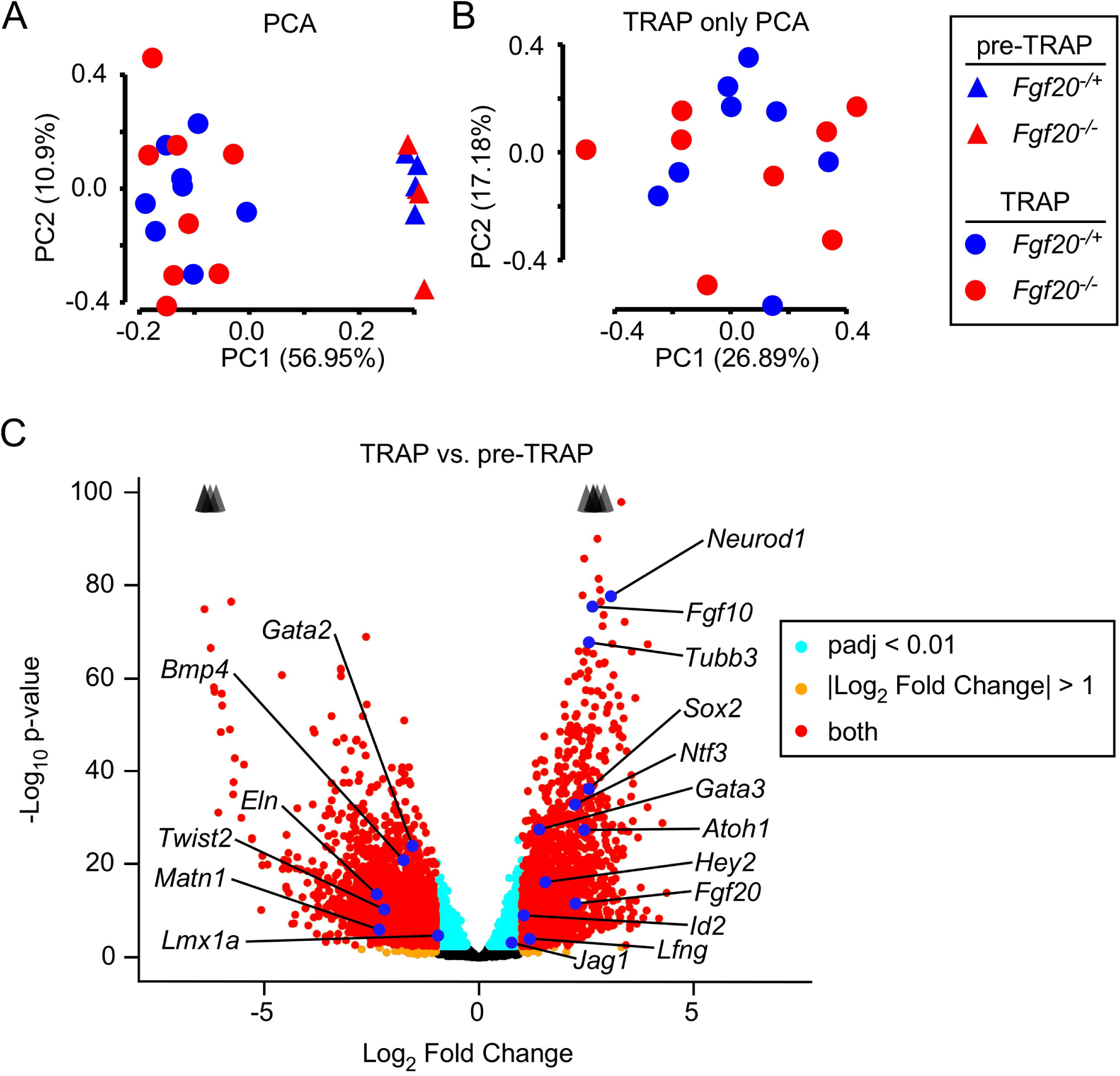
*Fgf20*^*Cre*^ TRAPseq enriched for prosensory domain mRNA. (A) Principal Component Analysis (PCA) on 24 TRAPseq samples (8 pre-TRAP samples – 4 *Fgf20*^*-/+*^, 4 *Fgf20*^*-/-*^; 16 TRAP samples – 8 *Fgf20*^*-/+*^, 8 *Fgf20*^*-/-*^) showing separation of pre-TRAP and TRAP samples along principal component (PC) 1, but not of *Fgf20*^*-/+*^ and *Fgf20*^*-/-*^ samples. (B) PCA on the 16 TRAP samples (excluding the 8 pre-TRAP samples) also did not show separation between *Fgf20*^*-/+*^ and *Fgf20*^*-/-*^ samples along the first two principal components. (C) Volcano plot showing TRAP vs. pre-TRAP differentially expressed genes. Positive Log_2_ Fold Change value indicates enrichment by TRAP; negative Log_2_ Fold Change value indicates depletion by TRAP. Labeled genes represent markers of the prosensory domain, Kölliker’s organ, spiral ganglion, outer sulcus, periotic mesenchyme, otic capsule. Padj, adjusted p-value for multiple comparisons (Benjamini-Hochberg method). The p-value plotted on y-axis is unadjusted. Arrowheads indicate genes above y-axis range.

### *Fgf20*^*Cre*^ TRAPseq revealed known FGF target genes during cochlear sensory epithelium differentiation

DEG analysis on *Fgf20*^*-/+*^ vs. *Fgf20*^*-/-*^ pre-TRAP samples resulted in, as expected, very few DEGs. In fact, only three genes were found to be significantly changed, based on adjusted p-value (padj) of < 0.1: *Tectb, Calb1*, and *Fgf20*. DEG analysis on *Fgf20*^*-/+*^ vs. *Fgf20*^*-/-*^ TRAP samples resulted in 47 DEGs with padj < 0.01 and 104 DEGs with padj < 0.1 (Fig. 3A). GO analysis with the top 362 TRAPseq DEGs (cut-off of padj < 0.5) found among the top 40 terms “sensory perception of sound,” “sensory organ morphogenesis,” “ear development,” and “inner ear receptor cell differentiation” (Table 3). Many neuronal and cell cycle biological processes terms, such as “regulation of neuron differentiation,” “forebrain neuron differentiation,” “regulation of neural precursor cell proliferation,” “cell division,” and “cell cycle arrest” were also among the top terms. *Fgf20*^*-/+*^ vs. *Fgf20*^*-/-*^ sequencing comparison data are presented in Supplemental file S2.

**Table 3.** Top enriched gene ontology (GO) terms from a list of top 362 *Fgf20*^*-/+*^ vs. *Fgf20*^*-/-*^ differentially expressed genes.

| Rank* | GO ID | GO biological processes term | p-value |
| --- | --- | --- | --- |
| 1 | GO:0003184 | pulmonary valve morphogenesis | 5.40E-06 |
| 2 | GO:0045664 | regulation of neuron differentiation | 6.80E-05 |
| 3 | GO:0007605 | sensory perception of sound | 1.50E-04 |
| 5 | GO:0007601 | visual perception | 3.60E-04 |
| 6 | GO:0001709 | cell fate determination | 4.10E-04 |
| 7 | GO:0046426 | negative regulation of JAK-STAT cascade | 4.10E-04 |
| 9 | GO:0021879 | forebrain neuron differentiation | 6.10E-04 |
| 12 | GO:0051301 | cell division | 1.12E-03 |
| 13 | GO:0021795 | cerebral cortex cell migration | 1.21E-03 |
| 14 | GO:0009948 | anterior/posterior axis specification | 1.23E-03 |
| 21 | GO:0090596 | sensory organ morphogenesis | 1.77E-03 |
| 28 | GO:0043583 | ear development | 2.50E-03 |
| 30 | GO:2000177 | regulation of neural precursor cell proliferation | 2.70E-03 |
| 33 | GO:0045596 | negative regulation of cell differentiation | 3.29E-03 |
| 35 | GO:0031175 | neuron projection development | 3.46E-03 |
| 38 | GO:0060113 | inner ear receptor cell differentiation | 4.33E-03 |
| 39 | GO:0007050 | cell cycle arrest | 4.76E-03 |

\* Rank by p-value (lowest to highest).

**Figure 3.**
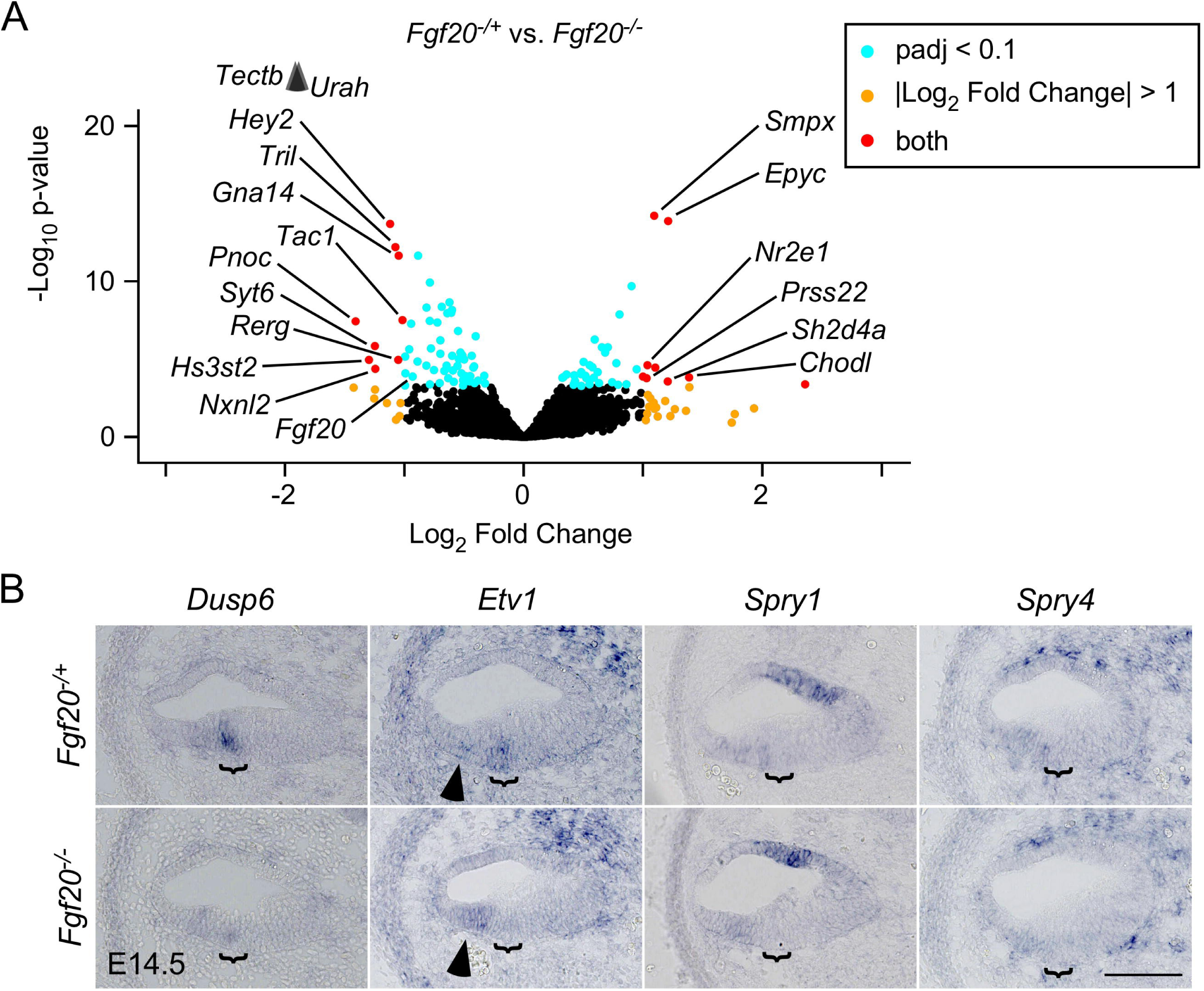
*Fgf20*^*Cre*^ TRAPseq revealed known FGF target genes during cochlear sensory epithelium differentiation. (A) Volcano plot showing *Fgf20*^*-/+*^ vs. *Fgf20*^*-/-*^ differentially expressed genes. *Fgf20* and transcripts meeting the criteria padj < 0.1 and Log_2_ Fold Change < −1 or > 1 are labeled, except predicted genes and unnamed transcripts. padj, adjusted p-value for multiple comparisons (Benjamini-Hochberg method). The p-value plotted on y-axis is unadjusted. Arrowheads indicate genes above y-axis range. (B) RNA in situ hybridization for known FGF target genes *Dusp6, Etv1, Spry1*, and *Spry4* on sections through the middle turn of E14.5 *Fgf20*^*-/+*^ (*Fgf20*^*Cre/+*^) and *Fgf20*^*-/-*^ (*Fgf20*^*Cre/βgal*^) cochlear ducts. Bracket, prosensory domain. Arrowhead, increased expression of *Etv1* in the outer sulcus of *Fgf20*^*-/-*^ cochleae. Scale bar, 100 µm.

For DEG analysis, we considered padj < 0.1 to be statistically significant. Confirming the validity of our TRAPseq results, DEGs with padj < 0.1 include *Fgf20* as well as *Hey1, Hey2, Etv4*, and *Etv5* (Table 4), which we have previously shown by RNA in situ hybridization (ISH) are downregulated in *Fgf20*^*-/-*^ vs. *Fgf20*^*-/+*^ cochleae (Yang et al. 2019). To confirm other genes identified by TRAPseq, we examined their expression patterns via ISH in *Fgf20*^*-/+*^ (*Fgf20*^*Cre/+*^) and *Fgf20*^*-/-*^ (*Fgf20*^*Cre/βgal*^) E14.5 cochleae. We began with DEGs that have been well-linked to FGF signaling (Table 4) and were downregulated in *Fgf20*^*-/-*^ cochleae by TRAPseq, such as *Dusp6, Etv1, Spry1*, and *Spry4* (Minowada et al. 1999; Münchberg and Steinbeisser 1999; Willardsen et al. 2014; Yang et al. 2018). By ISH, *Dusp6* (Dual specificity phosphatase 6) was expressed within the prosensory domain in control cochleae, and was almost undetectable in *Fgf20*^*-/-*^ cochleae (Fig. 3B). *Etv1* (Ets variant 1) was also expressed within the prosensory domain. Interestingly, while *Etv1* expression was absent in the prosensory domain in *Fgf20*^*-/-*^ cochleae, it was increased in the outer sulcus (Fig. 3B, arrowhead). *Spry1* (Sprouty homolog 1) and *Spry4* (Sprouty homolog 4) expressions were found diffusely in the floor of the cochlear duct, and appeared slightly decreased in *Fgf20*^*-/-*^ cochleae, although it was difficult to tell definitively by ISH (Fig. 3B).

**Table 4.** *Fgf20*^*-/+*^ vs. *Fgf20*^*-/-*^ differentially expressed genes associated with FGF signaling.

| Rank* | Ensembl ID | Gene | Enrichment<sup>^</sup> | Log2FC<sup>&</sup> | padj<sup>#</sup> |
| --- | --- | --- | --- | --- | --- |
| 9 | ENSMUSG000000019960 | *Dusp6* | enriched | -0.79 | <0.001 |
| 21 | ENSMUSG000000004151 | *Etv1* | depleted | -0.72 | <0.001 |
| 36 | ENSMUSG000000017724 | *Etv4* | ENRICHED | -0.60 | <0.01 |
| 23 | ENSMUSG000000013089 | *Etv5* | - | -0.55 | <0.001 |
| 74 | ENSMUSG000000031603 | *Fgf20* | ENRICHED | -0.93 | 0.03 |
| 50 | ENSMUSG000000040289 | *Hey1* | ENRICHED | -0.55 | 0.01 |
| 5 | ENSMUSG000000019789 | *Hey2* | ENRICHED | -1.12 | <0.001 |
| 106 | ENSMUSG000000037211 | *Spry1* | ENRICHED | -0.45 | 0.10 |
| 87 | ENSMUSG000000024427 | *Spry4* | depleted | -0.45 | 0.06 |

\* Rank by padj (lowest to highest).
<sup>^</sup> Enrichment by TRAP: results of TRAP vs. pre-TRAP comparison. Enriched indicates Log<sub>2</sub> Fold Change > 0 and padj < 0.05. Depleted indicates Log<sub>2</sub> Fold Change < 0 and padj < 0.05. Upper case indicates Log<sub>2</sub> Fold Change > 1 or < -1. Dash (-) indicates padj > 0.05.
<sup>&</sup> Log<sub>2</sub> Fold Change of *Fgf20*<sup>+/+</sup> vs. *Fgf20*<sup>-/-</sup> comparison.
<sup>#</sup> Adjusted p-value of *Fgf20*<sup>+/+</sup> vs. *Fgf20*<sup>-/-</sup> comparison.

### *Fgf20*^*Cre*^ TRAPseq revealed many genes associated with cochlea development or hearing loss

Many DEGs from *Fgf20*^*-/+*^ vs. *Fgf20*^*-/-*^ TRAPseq have previously been associated with cochlea development (Table 5). We validated some interesting DEGs via ISH, including *Tectb, Smpx, Epyc, Fat3*, and *Heyl* (Fig. 4A). *Tectb* (Tectorin beta) was expressed in the prosensory domain and Kölliker’s organ and was nearly absent in the prosensory domain of *Fgf20*^*-/-*^ cochleae. Meanwhile, *Tecta* (Tectorin alpha), which also trended towards lower expression per TRAPseq (padj = 0.22), was not changed based on ISH. *Smpx* (Small muscle protein, X-linked) was lowly expressed in the prosensory domain and was increased in *Fgf20*^*-/-*^ cochleae. *Epyc* (Epiphycan) was faintly expressed in the medial cochlear duct wall at this stage and was increased in *Fgf20*^*-/-*^ cochleae. *Fat3* (FAT atypical cadherin 3) was expressed in the prosensory domain and was decreased in *Fgf20*^*-/-*^ cochleae. *Heyl* (hairy/enhancer-of-split related with YRPW motif-like) has not been associated with cochlea development or hearing loss, but belongs in the same family as *Hey1* and *Hey2*. By ISH, it is not expressed in the cochlea at E14.5, but is upregulated in the prosensory domain in *Fgf20*^*-/-*^ cochleae.

**Table 5.** *Fgf20*^*-/+*^ vs. *Fgf20*^*-/-*^ differentially expressed genes associated with hearing or cochlear development.

| Rank* | Ensembl ID | Gene | Enrichment^ | Log2FC& | padj# |
| --- | --- | --- | --- | --- | --- |
| 236 | ENSMUSG000000021835 | *Bmp4* | DEPLETED | 0.41 | 0.38 |
| 10 | ENSMUSG000000028222 | *Calb1* | ENRICHED | 0.90 | <0.001 |
| 16 | ENSMUSG000000027555 | *Car13* | enriched | -0.64 | <0.001 |
| 331 | ENSMUSG00000003031 | *Cdkn1b* | - | -0.43 | 0.47 |
| 11 | ENSMUSG000000030862 | *Cpxm2* | enriched | -0.62 | <0.001 |
| 12 | ENSMUSG000000030905 | *Crym* | depleted | -0.69 | <0.001 |
| 182 | ENSMUSG000000043969 | *Emx2* | - | -0.61 | 0.27 |
| 218 | ENSMUSG000000087095 | *Emx2os* | - | -0.60 | 0.35 |
| 4 | ENSMUSG000000019936 | *Epyc* | depleted | 1.21 | <0.001 |
| 31 | ENSMUSG000000074505 | *Fat3* | - | -0.96 | <0.01 |
| 47 | ENSMUSG000000015053 | *Gata2* | DEPLETED | 0.55 | <0.01 |
| 28 | ENSMUSG000000032744 | *Heyl* | DEPLETED | 0.65 | <0.01 |
| 119 | ENSMUSG000000050100 | *Hmx2* | - | 0.74 | 0.11 |
| 96 | ENSMUSG000000026686 | *Lmx1a* | depleted | 0.55 | 0.08 |
| 49 | ENSMUSG000000098318 | *Lockd* | - | -0.65 | 0.01 |
| 29 | ENSMUSG000000036446 | *Lum* | - | 0.70 | <0.01 |
| 61 | ENSMUSG000000027210 | *Meis2* | DEPLETED | 0.48 | 0.02 |
| 168 | ENSMUSG000000030739 | *Myh14* | ENRICHED | -0.41 | 0.23 |
| 97 | ENSMUSG000000004891 | *Nes* | DEPLETED | -0.78 | 0.08 |
| 83 | ENSMUSG000000060424 | *Pantr1* | depleted | -0.51 | 0.05 |
| 53 | ENSMUSG000000045515 | *Pou3f3* | depleted | -0.42 | 0.01 |
| 71 | ENSMUSG000000031665 | *Sall1* | enriched | -0.49 | 0.03 |
| 177 | ENSMUSG000000049532 | *Sall2* | ENRICHED | -0.39 | 0.26 |
| 89 | ENSMUSG000000024565 | *Sall3* | enriched | -0.59 | 0.06 |
| 14 | ENSMUSG000000035109 | *Shc4* | ENRICHED | -0.60 | <0.001 |
| 3 | ENSMUSG000000041476 | *Smpx* | ENRICHED | 1.09 | <0.001 |
| 225 | ENSMUSG000000074637 | *Sox2* | ENRICHED | -0.45 | 0.37 |
| 18 | ENSMUSG000000061762 | *Tac1* | - | -1.01 | <0.001 |
| 161 | ENSMUSG000000037705 | *Tecta* | ENRICHED | -0.36 | 0.22 |
| 1 | ENSMUSG000000024979 | *Tectb* | ENRICHED | -1.91 | <0.001 |
| 128 | ENSMUSG000000021779 | *Thrb* | ENRICHED | 0.45 | 0.13 |

\* Rank by padj (lowest to highest).
^ Enrichment by TRAP: results of TRAP vs. pre-TRAP comparison. Enriched indicates Log<sub>2</sub> Fold Change > 0 and padj < 0.05. Depleted indicates Log<sub>2</sub> Fold Change < 0 and padj < 0.05. Upper case indicates Log<sub>2</sub> Fold Change > 1 or < -1. Dash (-) indicates padj > 0.05.
& Log<sub>2</sub> Fold Change of *Fgf20*<sup>+/+</sup> vs. *Fgf20*<sup>-/-</sup> comparison.

**Figure 4.**
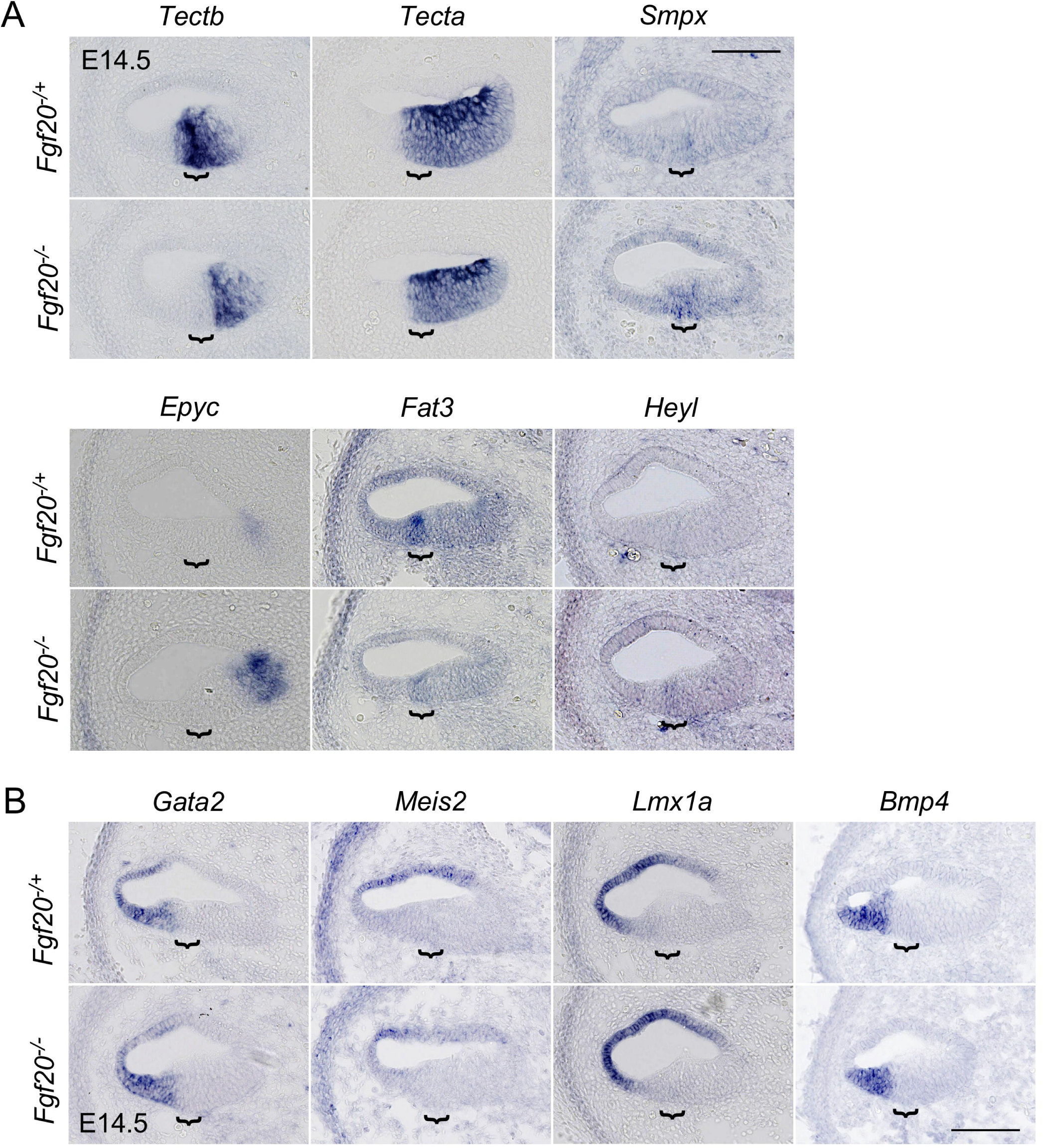
*Fgf20*^*Cre*^ TRAPseq revealed many genes associated with cochlea development or hearing loss. RNA in situ hybridization on sections through the middle turn of E14.5 *Fgf20*^*-/+*^ (*Fgf20*^*Cre/+*^) and *Fgf20*^*-/-*^ (*Fgf20*^*Cre/βgal*^) cochlear ducts. Bracket, prosensory domain. Scale bar, 100 µm. (A)Genes *Tectb, Tecta, Smpx, Epyc, Fat3*, and *Heyl* (B) Genes *Gata2, Meis2, Lmx1a*, and *Bmp4*

TRAPseq also identified a few transcription factors that were depleted by TRAP, but increased in *Fgf20*^*-/-*^ cochleae, including *Gata2* (GATA binding protein 2*), Meis2* (Meis homeobox 2), and *Lmx1a* (LIM homeobox transcription factor 1 alpha). Depletion by TRAP suggests that they are not highly expressed in the prosensory domain or Kölliker’s organ. By ISH, all three genes were expressed in the outer sulcus and/or roof of the cochlear duct (Fig. 4B). However, ISH did not appear to be sensitive enough to detect differences in the expression of any of these genes. *Bmp4* (Bone morphogenetic protein 4) is another gene depleted by TRAP, but not significantly changed in *Fgf20*^*-/-*^ cochleae (padj = 0.38). By ISH *Bmp4* was expressed in the outer sulcus and did not show any changes in *Fgf20*^*-/-*^ cochleae (Fig. 4B).

### *Fgf20*^*Cre*^ TRAPseq revealed decreased expression of cell cycle regulators

GO analysis on *Fgf20*^*-/+*^ vs. *Fgf20*^*-/-*^ TRAPseq DEGs showed that the cell cycle may be affected by the loss of FGF20, with the terms “cell division” and “cell cycle arrest” among the top terms (Table 3). This was confirmed by known and predicted protein-protein interaction (PPI) network identification via the STRING database (Snel et al. 2000; Szklarczyk et al. 2019) with the top 192 DEGs, representing those with padj < 0.3. By far the largest PPI network identified consisted of cell cycle regulators (Fig. 5A). The list of top DEGs indeed showed many genes involved in cell cycle regulation, all of which were decreased in expression in *Fgf20*^*-/-*^ cochleae (Table 6).

**Table 6.** *Fgf20*^*-/+*^ vs. *Fgf20*^*-/-*^ differentially expressed genes associated with cell cycle regulation.

| Rank* | Ensembl ID | Gene | Enrichment^ | Log2FC& | padj# |
| --- | --- | --- | --- | --- | --- |
| 70 | ENSMUSG000000027715 | *Ccna2* | depleted | -0.33 | 0.03 |
| 32 | ENSMUSG000000070348 | *Ccnd1* | - | -0.53 | <0.01 |
| 120 | ENSMUSG000000033102 | *Cdc14b* | - | -0.49 | 0.11 |
| 24 | ENSMUSG000000006398 | *Cdc20* | depleted | -0.40 | <0.001 |
| 222 | ENSMUSG000000024791 | *Cdca5* | - | -0.30 | 0.37 |
| 183 | ENSMUSG000000028873 | *Cdca8* | - | -0.33 | 0.28 |
| 197 | ENSMUSG000000019942 | *Cdk1* | depleted | -0.29 | 0.31 |
| 206 | ENSMUSG000000026023 | *Cdk15* | ENRICHED | -0.45 | 0.32 |
| 121 | ENSMUSG000000037628 | *Cdkn3* | depleted | -0.41 | 0.12 |
| 159 | ENSMUSG000000026605 | *Cenpf* | - | -0.92 | 0.22 |
| 158 | ENSMUSG000000001517 | *Foxm1* | depleted | -0.43 | 0.22 |
| 95 | ENSMUSG000000027331 | *Knstrn* | - | -0.32 | 0.07 |
| 146 | ENSMUSG000000020808 | *Pimreg* | - | -0.37 | 0.18 |
| 147 | ENSMUSG000000030867 | *Plk1* | depleted | -0.38 | 0.18 |

\* Rank by padj (lowest to highest).
^ Enrichment by TRAP: results of TRAP vs. pre-TRAP comparison. Enriched indicates Log<sub>2</sub> Fold Change > 0 and padj < 0.05. Depleted indicates Log<sub>2</sub> Fold Change < 0 and padj < 0.05. Upper case indicates Log<sub>2</sub> Fold Change > 1 or < -1. Dash (-) indicates padj > 0.05.
& Log<sub>2</sub> Fold Change of *Fgf20*<sup>+/-</sup> vs. *Fgf20*<sup>-/-</sup> comparison.

**Figure 5.**
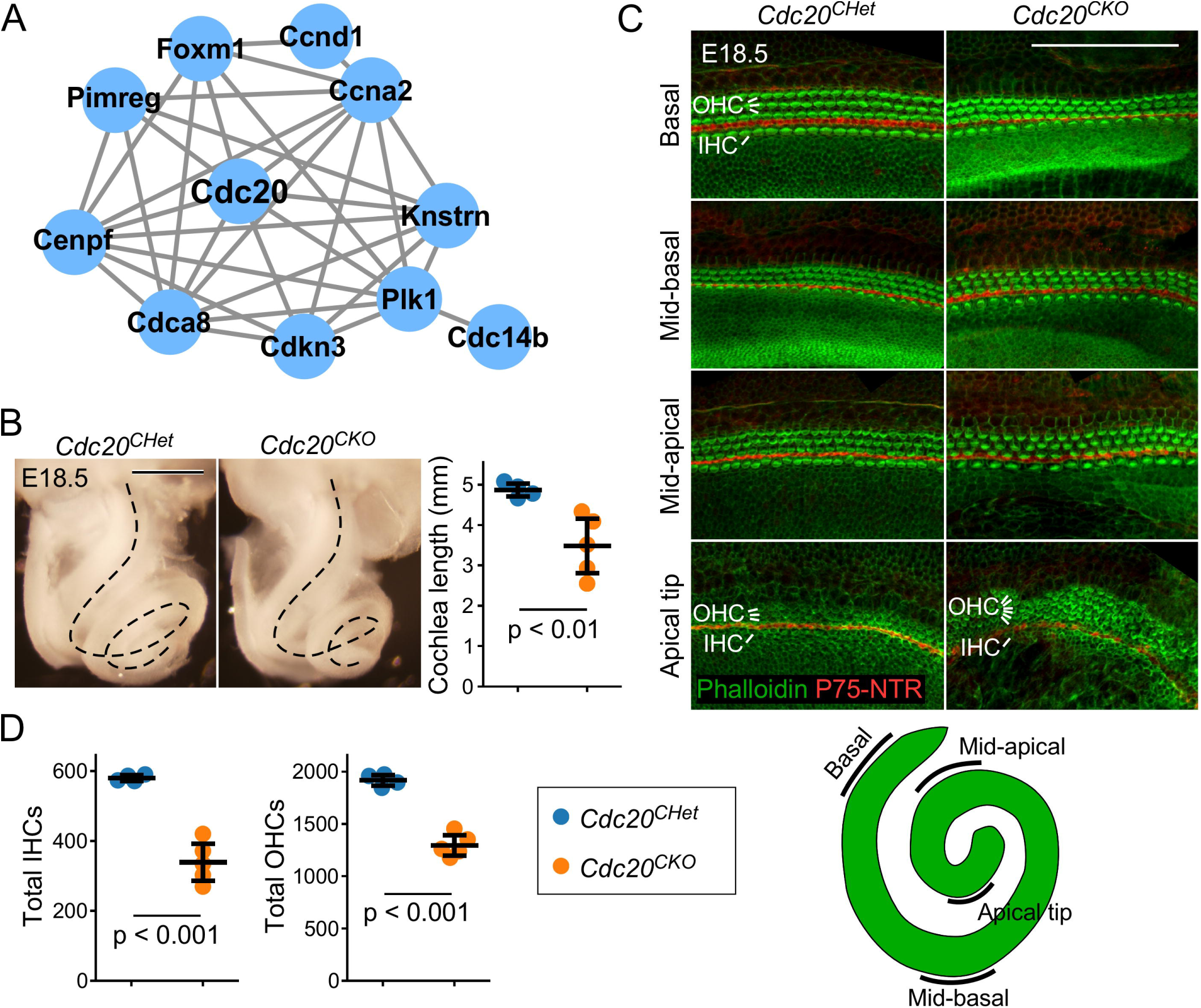
*Fgf20*^*Cre*^ TRAPseq revealed decreased expression of cell cycle regulators. (A) The largest protein-protein interaction network identified via the STRING database consisted of genes involved in cell cycle regulation. Lines represent known and predicted protein-protein interactions of high or very high confidence (minimum required interaction score = 0.700). (B) Dissected inner ears from E18.5 *Cdc20*^*CHet*^ (*Fgf20*^*Cre/+*^;*Cdc20*^*flox/+*^) and *Cdc20*^*CKO*^ (*Fgf20*^*Cre/+*^;*Cdc20*^*flox/flox*^) embryos with otic capsule removed to reveal the cochlea (dotted lines). Scale bar, 0.5 mm. Quantification of cochlea length measured using whole mount cochlea *Cdc20*^*CHet*^ (n = 4) and *Cdc20*^*CKO*^ (n = 5). Error bars represent mean ± std. Results were analyzed by Student’s t-test; p-values are shown. (C) Whole mount cochlea from E18.5 *Cdc20*^*CHet*^ and *Cdc20*^*CKO*^ embryos showing one row of inner hair cells (IHC) and three rows of outer hair cells (OHC) marked by phalloidin (green) and separated by inner pillar cells (p75NTR, red). Representative regions from the basal, mid-basal, and mid-apical turns, and apical tip of the cochlea are shown. See schematic below showing locations of the turns of the cochlea. At the apical tip, four or more rows of OHCs frequently observed in *Cdc20*^*CHet*^ cochleae. Scale bar, 100 µm. (D) Quantification of total number of inner and outer hair cells (IHCs and OHCs) in E18.5 *Cdc20*^*CHet*^ (n = 4) and *Cdc20*^*CKO*^ (n = 5) cochleae. Error bars represent mean ± std. Results were analyzed by Student’s t-test; p-values are shown.

Although we have not found that *Fgf20* regulates cell cycle progression by itself, *Fgf20* does interact with *Fgf9* and *Sox2* to regulate prosensory progenitor and Kölliker’s organ proliferation, respectively (Huh et al. 2015; Yang et al. 2019). We hypothesize, therefore, that the expression changes in cell cycle regulators may reflect these functions of *Fgf20*. To rule out the possibility that cell cycle regulation contributes to the differentiation and patterning defect found in *Fgf20*^*-/-*^ cochleae, we examined the largest node of the PPI network, *Cdc20* (Cell division cycle 20). *Cdc20* is a coactivator of the anaphase-promoting complex (APC), the cell cycle-regulated ubiquitin ligase. Interestingly, Cdc20-APC is required for presynaptic axon differentiation in postmitotic neurons in the cerebellum (Yang et al. 2009).

To examine how decreased expression of *Cdc20* may contribute to the *Fgf20*^*-/-*^ phenotype, we combined *Fgf20*^*Cre*^ with *Cdc20*^*flox*^ to conditionally delete *Cdc20* from the *Fgf20*^*Cre*^ lineage (Manchado et al. 2010). *Fgf20*^*Cre/+*^;*Cdc20*^*flox/flox*^ (*Cdc20*^*CKO*^) cochleae (length: 3.48 ± 0.75 mm) were shorter and more tightly coiled than *Fgf20*^*Cre/+*^;*Cdc20*^*flox/+*^ (*Cdc20*^*CHet*^) cochleae (length: 4.86 ± 0.18 mm) (Fig. 5B). Importantly, HCs (labeled by phalloidin, green) in *Cdc20*^*CKO*^ cochleae exhibited the normal pattern of one row of IHCs separated from three rows of OHCs by pillar cells (inner pillar cells labeled by P75^NTR^, red) (Fig. 5C). Interestingly, 4 of 5 *Cdc20*^*CKO*^ cochleae examined had 4 or more rows of OHCs at the apical tip (Fig. 5C), which may be the result of a defect in convergent extension. Upon quantification, total number of IHCs and OHCs were decreased in *Cdc20*^*CKO*^ (IHC: 339 ± 59; OHC: 1295 ± 108) relative to *Cdc20*^*CHet*^ (IHC: 580 ± 9; OHC: 1916 ± 59) cochleae (Fig. 5D); however, this can likely be attributed to the shorter length of the cochlea.

### *Sall1*-ΔZn^2-10^ mutant cochleae exhibit an outer hair cell phenotype

*Sall1* (Sal-like 1) and *Sall3* (Sal-like 3), members of a family of transcription factors, are expressed in the cochlear duct throughout development (Nishinakamura et al. 2001; Ott et al. 1996, 2001; Parrish et al. 2004). Both were identified by TRAPseq as decreased in *Fgf20*^*-/-*^ cochleae (Table 5). ISH showed that both *Sall1* and *Sall3* were expressed in the prosensory domain and were decreased in *Fgf20*^*-/-*^ cochleae (Fig. 6A). Interestingly, *Sall2* (Sal-like 2), another member of the same family, trended towards lower expression according to TRAPseq (padj = 0.26). By ISH, *Sall2* was also expressed in the prosensory domain, but was not noticeably decreased in *Fgf20*^*-/-*^ cochleae (Fig. 6A). *Sall4*, the fourth member of the *Sall* family, was filtered out from TRAPseq analysis due to insufficiently low read counts.

**Figure 6.**
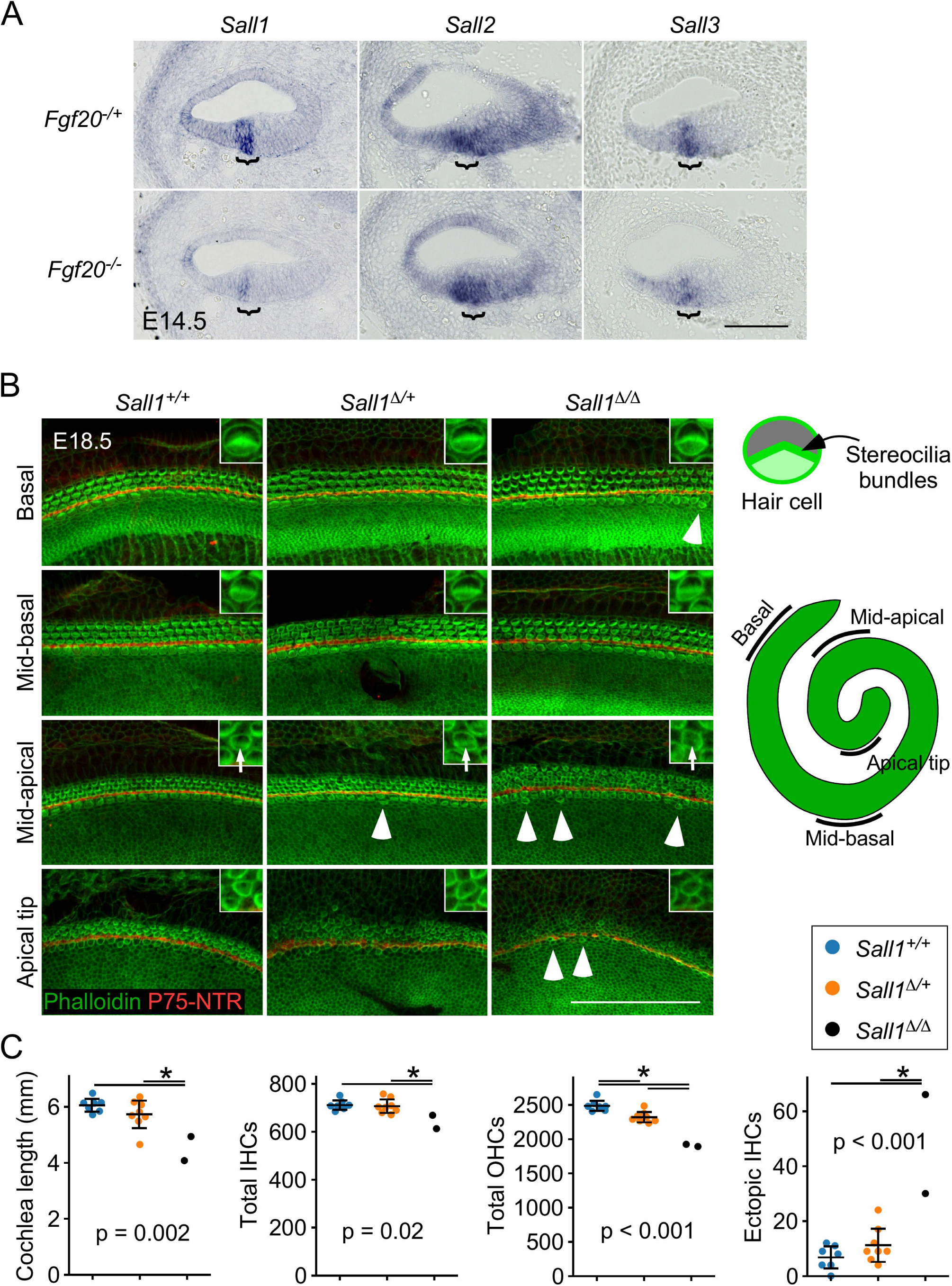
*Sall1-ΔZn2-10* mutant cochleae exhibit an outer hair cell phenotype. (A) RNA in situ hybridization for *Sall1, Sall2*, and *Sall3* on sections through the middle turn of E14.5 *Fgf20*^*-/+*^ (*Fgf20*^*Cre/+*^) and *Fgf20*^*-/-*^ (*Fgf20*^*Cre/βgal*^) cochlear ducts. Bracket, prosensory domain. Scale bar, 100 µm. (B) Whole mount cochlea from E18.5 *Sall1*^*+/+*^, *Sall1*^*Δ/+*^, and *Sall1*^*Δ/Δ*^ embryos showing inner hair cells and outer hair cells marked by phalloidin (green) and separated by inner pillar cells (p75NTR, red). Representative regions from the basal (5% of total length from the basal tip), mid-basal (33%), and mid-apical (67%) turns, and apical tip (90%) of the cochlea are shown. See schematic to the right showing locations of the turns of the cochlea. Inset: 3.8x magnified image of a representative OHC showing stereocilia bundle formation (arrows in mid-apical region). Numerous ectopic inner hair cells were found throughout the *Sall1*^*Δ/Δ*^ cochleae, especially towards the apex (arrowheads). Scale bar, 100 µm. (C) Quantification of cochlea length, total number of inner hair cells (IHCs) and outer hair cells (OHCs), and total number of ectopic IHCs in E18.5 *Sall1*^*+/+*^ (n = 7), *Sall1*^*Δ/+*^ (n = 8), and *Sall1*^*Δ/Δ*^ (n = 2). Error bars represent mean ± std. Results were analyzed by one-way ANOVA. P-values shown are from the ANOVA. * indicates p < 0.05 from Tukey’s HSD (ANOVA post-hoc).**Table 1**. Top 12 enriched gene ontology (GO) terms from a list of 2017 differentially expressed genes depleted by TRAP, compared to pre-TRAP samples.

Importantly, *SALL1* has been linked to Townes-Brocks syndrome (TBS) in humans, which causes sensorineural hearing loss, among other developmental defects (Kohlhase et al. 1998). Mutations in one copy of *SALL1* is responsible for TBS, although *SALL1* haploinsufficiency may not be the sole causative factor, as *Sall1*-null mice do not recapitulate the human TBS phenotypes (Nishinakamura et al. 2001). Instead, mice expressing one copy of a *Sall1* allele with mutations known to cause TBS, *Sall1-ΔZn*^*2-10*^ (*Sall1*^*Δ*^), mimic TBS defects, including hearing loss (Kiefer et al. 2003). This mutation results in a truncated protein encoding the N-terminus of Sall1, which has been shown to mediate transcriptional repression (Kiefer et al. 2002). Like wildtype Sall1, the truncated Sall1^Δ^ protein can bind all members of the Sall family (Kiefer et al. 2003), and its expression alone in transgenic mice leads to derepression of Sall-regulated genes resulting in TBS-like phenotypes (Kiefer et al. 2008). These results suggest that Sall1^Δ^ may act as a dominant negative and interfere with the transcription-repressor activity of all Sall proteins.

We hypothesized that the dominant negative effects of Sall^Δ^ may recapture the decrease in *Sall1* and *Sall3* expression in *Fgf20*^*-/-*^ cochleae. To see if mice heterozygous for this mutation (*Sall1*^*Δ/+*^) may exhibit cochlea development phenotypes similar to *Fgf20*^*-/-*^ mice, we examined *Sall1*^*Δ/+*^ cochleae at E18.5. While the overall HC patterning appeared unchanged in *Sall1*^Δ/+^ cochleae compared to *Sall1*^*+/+*^ (Fig. 6B), there was a small but statistically significant decrease in the number of OHCs in *Sall1*^*Δ/+*^ cochleae (2321 ± 79), compared to *Sall1*^*+/+*^ (2486 ± 81) (Fig. 6C). The number of IHCs (*Sall1*^*+/+*^: 711 ± 21; *Sall1*^Δ/+^: 707 ± 30 mm) appeared unchanged, suggesting that OHCs may be more sensitive to the *Sall1*^Δ^ mutation. Cochlea length (*Sall1*^*+/+*^: 6.06 ± 0.25 mm; *Sall1*^Δ/+^: 5.73 ± 0.53 mm including a possible outlier at 4.65 mm) also appeared relatively unchanged.

Most embryos homozygous for the *Sall1*^*Δ*^ mutation (*Sall1*^*Δ/Δ*^) die by E16.5 (Kiefer et al. 2003). However, we were able to obtain two *Sall1*^Δ/Δ^ embryos that survived to E18.5. The cochleae of these embryos showed a further reduction of the number of OHCs compared to *Sall1*^*Δ/+*^ (1898 and 1922 in the two samples, a 23-24% decrease compared to *Sall1*^*+/+*^ cochleae). *Sall1*^Δ/Δ^ embryos also showed a decrease in the number of IHCs (612 and 668 in the two samples, a 9-14% decrease compared to *Sall1*^*+/+*^ cochleae) and in cochlea length (4.07 mm and 4.94 mm in the two samples, a 18-32% decrease compared to *Sall1*^*+/+*^ cochleae) (Fig. 6C). The decrease in the number of OHCs is more severe than the decrease in number of IHCs, again suggesting that OHCs may be more sensitive to the *Sall1*^*Δ*^ mutation.

In addition, the HCs in *Sall1*^*Δ/Δ*^ cochleae appeared less mature than those found in comparative regions of *Sall1*^*+/+*^ and *Sall1*^*Δ/+*^ cochleae, based on F-actin organization in phalloidin stained samples (Fig. 6B). This is most apparent in the mid-apical turns, where stereocilia bundles appeared relatively well-formed in *Sall1*^*+/+*^ and *Sall1*^*Δ/+*^ cochleae (Fig. 6B, inset). In *Sall1*^*Δ/Δ*^ cochleae, however, the stereocilia in this region appeared much more immature and disorganized (arrows in Fig. 6B inset), resembling those found in the less mature apical tip (hair cell differentiation and maturation occur in a base-to-apex gradient (Basch et al. 2016), therefore, more apical hair cells are less mature). In the apical tip, the F-actin networks at the HC cortex in *Sall1*^*Δ/Δ*^ cochleae appeared less dense than those found in *Sall1*^*+/+*^ and *Sall1*^*Δ/+*^ cochleae, as indicated by weaker phalloidin labeling (Fig. 6B, inset).

Interestingly, many ectopic IHCs were found throughout the length of *Sall1*^*Δ/Δ*^ cochleae, especially towards the apex (Fig. 6B, arrowheads). Quantification of these ectopic IHCs showed a statistically significant increase in *Sall1*^*Δ/Δ*^ (30 and 66) compared to *Sall1*^*+/+*^ (7 ± 4) and *Sall1*^*Δ/+*^ (11 ± 6) cochleae (Fig. 6C). These ectopic IHCs suggest a patterning defect in *Sall1*^*Δ/Δ*^ cochleae.

## DISCUSSION

We have adapted the TRAP technique to study a relatively small population of difficult-to-isolate cells: cochlear prosensory cells. TRAP using *Fgf20*^*Cre*^ combined with *ROSA*^*fsTRAP*^ effectively enriched for translating mRNA from the *Fgf20*^*Cre*^ lineage at E14.5. We believe the pre-TRAP vs. TRAP DEG analysis provides a useful dataset for identifying genes enriched in the prosensory domain, Kölliker’s organ, and spiral ganglion of the developing cochlea.

TRAPseq comparing *Fgf20*^*-/+*^ and *Fgf20*^*-/-*^ E14.5 cochlea samples showed decreased expression of known FGF20 signaling targets in the cochlea at this stage: *Etv4, Etv5, Hey1*, and *Hey2*. It also showed decreased expression of other FGF signaling targets, such as *Dusp6* and *Etv1*, further confirming the validity our technique. Just as importantly, TRAPseq DEGs did not include genes that we have previously shown are not downstream of FGF20, but that have been shown to be downstream of FGFR1: *Cdkn1b* and *Sox2* (Table 5) (Huh et al. 2015; Yang et al. 2019). Interestingly, however, *Lockd*, a non-coding RNA near the *Cdkn1b* locus and co-expressed with *Cdkn1b* (Paralkar et al. 2016), was significantly decreased in *Fgf20*^*-/-*^ cochleae (Table 5). The expression of a few other genes previously shown to be downstream of FGFR1 during cochlea development, such as *Fgf10, Hes5*, and *Ntf3* (Ono et al. 2014; Pirvola et al. 2002), were also not significantly changed.

As with any large data experiment, false positives and negatives are expected. Here, we used a lenient false discovery rate of 0.1 to evaluate *Fgf20*^*-/+*^ and *Fgf20*^*-/-*^ TRAPseq results to reduce the number of false negatives at the cost of increasing false positives. While we were able to confirm many TRAPseq DEGs via ISH, as well as confirm the expression of several non-significantly changed genes as unchanged via ISH, there were discrepancies between TRAPseq and ISH results. Besides false positivity, another possible and interesting explanation for the discrepancies is that TRAPseq specifically identifies differences in translating mRNA. Such differences may not always be reflected in the whole mRNA population detected by ISH, due to posttranscriptional regulation. Therefore, TRAPseq data may be a more accurate representation of protein expression.

Another caveat is that the *Fgf20*^*Cre*^-TRAP enrichment process is not perfect, due to limitations of the technique and the inclusion of Kölliker’s organ and spiral ganglion cells in the *Fgf20*^*Cre*^ lineage. RNA from these sources dilute the mRNA from the target prosensory cell population, reducing the power of TRAPseq in detecting changes within this population.

### TRAPseq identified DEGs previously associated with cochlea development or hearing

A few other DEGs identified by TRAPseq comparison of *Fgf20*^*-/+*^ and *Fgf20*^*-/-*^ cochleae have known roles in cochlea development (Table 5). Altered expression of these genes, therefore, 5 may contribute to the *Fgf20*^*-/-*^ phenotype. Importantly, we do not know what proportion of these DEGs are directly regulated by FGF20, and what proportion may be indirectly regulated or are markers of dysregulated differentiation. Here, we highlight some of these DEGs.

*Fat3*, encoding a mammalian homolog of the *Drosophila* cell adhesion molecule Fat, is required for the normal patterning of OHCs, along with *Fat4* (Saburi et al. 2012). *Fat3*-null cochleae exhibits a small loss of OHCs from the base of the cochlea and a slight gain of OHCs in the mid-apex. We hypothesize that the decreased expression of *Fat3* may contribute to the OHC patterning defect in *Fgf20*^*-/-*^ cochleae.

*Cpxm2* (Carboxypeptidase X 2) is one of the three genes on chromosome 7 deleted in the head bobber mouse line, which exhibits deafness and vestibular defects (Buniello et al. 2013; Somma et al. 2012). However, how *Cpxm2* deletion contributes to deafness in these mice has not been elucidated.

*Tectb* is expressed in the prosensory domain (Rau et al. 1999) and encodes a major glycoprotein in the tectorial membrane required for normal hearing (Russell et al. 2007). Interestingly, *Tecta*, another gene encoding a glycoprotein in the tectorial membrane, trended towards lower expression (not statistically significantly) and did not show decreased expression by ISH. The composition of the tectorial membrane in *Fgf20*^*-/-*^ cochleae has not been studied.

*Thrb* (Thyroid hormone receptor beta) is expressed in the OC, GER, spiral ligament, and spiral ganglion in the neonatal cochlea. *Thrb*-mutant mice (both null and mutants with disrupted thyroid hormone binding) have severe hearing loss attributed to disruption of postnatal morphogenesis of the tectorial membrane (Forrest et al. 1996; Griffith et al. 2002; Kaukua et al. 2014; Ng et al. 2015; Sharlin et al. 2011). Interestingly, *Thrb* trended towards increased expression per TRAPseq (padj = 0.13).

*Myh14* (myosin, heavy polypeptide 14) is one of the genes encoding Myosin II. It is expressed in both developing HCs and SCs in the prenatal organ of Corti. Myosin II is required for patterning and convergent extension in the cochlea (Yamamoto et al. 2009). A convergence and extension defect may contribute to the shortened length of *Fgf20*^*-/-*^ cochleae. *Myh14* trended towards decreased expression per TRAPseq (padj = 0.23).

*Smpx*, previously shown to be expressed in HCs (Yoon et al. 2011), is associated with heritable progressive hearing loss (Abdelfatah et al. 2013; Huebner et al. 2011; Schraders et al. 2011). However, *Smpx*-null mice have not been shown to have a hearing defect or much of an overt developmental phenotype (Palmer et al. 2001). Given that *Smpx* is expressed in HCs, its increased expression in *Fgf20*^*-/-*^ cochleae may reflect the premature onset of HC differentiation in these mice.

*Epyc*, encoding a proteoglycan expressed in mature nonsensory regions of the cochlear duct, is required for normal hearing (Hanada et al. 2017). Its faint expression at E14.5 in the medial cochlear duct wall of control cochlea and increased expression in *Fgf20*^*-/-*^ cochleae may also reflect the premature onset of differentiation, although it has not been shown that the Kölliker’s organ undergoes premature differentiation in *Fgf20*^*-/-*^ cochleae.

### TRAPseq identified DEGs with unknown functions in cochlea development

Most of the DEGs identified by *Fgf20*^*-/+*^ and *Fgf20*^*-/-*^ TRAPseq have no known roles in cochlea development. However, some of these are related to genes with known roles in cochlea development, suggesting possible redundancy. Here, we highlight some of the most interesting ones.

*Dusp6* is a known downstream target of FGF signaling (Dickinson and Keyse 2006) and is a downstream target of FGF20 signaling in the olfactory system (Yang et al. 2018). Mice heterozygous for a *Dusp6*-null allele exhibit hearing loss, attributed to malformed otic capsule and ossicles (Li et al. 2007). While *Dusp6* is known to be expressed in the prosensory domain and the organ of Corti (Urness et al. 2008), which we confirm, its role in the development of these structures has not been investigated.

*Etv4* (Ets variant 4) and *Etv5* (Ets variant 5) have been shown to be downstream of FGF20/FGFR1 signaling in the developing cochlea (Hayashi et al. 2008; Yang et al. 2019). However, *Etv1*, the third member of the PEA3 group of Ets transcription factors, has not been associated with cochlea development. We show here that *Etv1* expression is decreased in the prosensory domain in *Fgf20*^*-/-*^ cochleae, while its expression is increased in the outer sulcus. This is potentially a significant pattern change, as the *Fgf20*^*-/-*^ phenotype is more severe in the outer compartment. Investigating whether this increase in expression in the outer sulcus contributes to the *Fgf20*^*-/-*^ phenotype will be addressed in future experiments.

*Hey1* and *Hey2* (hairy/enhancer-of-split related with YRPW motif 1 and 2) have been shown to be downstream of FGF20 signaling in the developing cochlea and are required to prevent premature HC differentiation (Benito-Gonzalez and Doetzlhofer 2014; Yang et al. 2019). TRAPseq identified that a third member of the Hes-related gene family, *Heyl*, is significantly increased in *Fgf20*^*-/-*^ cochleae at E14.5. Based on this observation, we hypothesize *Heyl* may be the compensating for the loss of *Hey1* and *Hey2*.

Other DEGs with unclear functional significance but that are known to be expressed in the cochlea include (Table 5):

- *Pou3f3* (POU domain, class 3, transcription factor 3) is expressed in SCs and mesenchymal cells in the cochlea (Mutai et al. 2009). Based on ISH from the Eurexpress atlas, it is also expressed in the cochlear duct floor at E14.5 (Diez-Roux et al., 2011, http://www.eurexpress.org euxassay_019559). However, analysis of the *Pou3f3*-null mouse cochlea did not reveal any apparent developmental defects (Mutai et al. 2009). Despite this, auditory and vestibular impairments have been reported in a *Pou3f3* (*Pou3f3*^*L423P*^) mutant mouse line (Kumar et al. 2016). Interestingly, *Pantr1* (Pou3f3 adjacent noncoding transcript 1), a lncRNC that shares a bidirectional promoter with *Pou3f3* (Goff et al. 2015), was also decreased in *Fgf20*^*-/-*^ cochleae per TRAPseq, suggesting disrupted activity at the promoter. In addition, *Rorb* (RAR-related orphan receptor beta), found to be increased in *Fgf20*^*-/-*^ cochleae by TRAPseq, has an antagonistic interaction with *Pou3f3* during cell fate specification in the developing neocortex (Oishi et al. 2016).
- *Calb1* (Calindin 1): expressed in mature HCs (Waldhaus et al. 2015). Upregulation may represent premature onset of HC differentiation in *Fgf20*^*-/-*^ cochleae.
- *Crym* (Crystallin, mu): a thyroid hormone binding protein, highly expressed in nonsensory regions of the cochlea in adult rats (Usami et al. 2008).
- *Shc4* (SHC family, member 4): an adaptor protein expressed in the cochlear duct floor at 34 E14.5 and E15.5 (Hawley et al. 2011).
- *Car13* (Carbonic anhydrase 13): expressed in nonsensory regions of the cochlea and the mesenchyme at E15.5 and neonatal stages (Wu et al. 2013)
- *Tac1* (Tachykinin 1): reported to be expressed in the cochlear epithelium during development (Radde-Gallwitz et al. 2004).
- *Lum* (Lumican): expressed in the otic capsule, some mesenchyme, and nonsensory regions of the cochlear duct (Ficker et al. 2004).
- *Nes* (Nestin): expressed in the spiral ganglion and parts of the prosensory domain at E14.5 and E15.5, as well as in mature SCs (Chow et al. 2016, 2015).

### Nonsensory cell markers are upregulated in *Fgf20*^*-/-*^ cochleae

*Fgf20*^*-/+*^ vs. *Fgf20*^*-/-*^ TRAPseq also identified a few transcription factors expressed in the outer sulcus and other nonsensory cochlear epithelium: *Gata2, Meis2*, and *Lmx1a* (Haugas et al. 2010; Koo et al. 2009; Lilleväli et al. 2004; Mann et al. 2017; Nichols et al. 2008; Sánchez-Guardado et al. 2011). As expected, all of these genes were depleted by TRAP, as they are not expressed in the prosensory domain or Kölliker’s organ. Interestingly, they are all increased in *Fgf20*^*-/-*^ cochleae per TRAPseq, suggesting that undifferentiated progenitors in *Fgf20*^*-/-*^ cochlea may have adopted a nonsensory identity. Two other outer sulcus/nonsensory epithelial markers, *Hmx2* and *Bmp4* (Morsli et al. 1998; Wang et al. 2001), also trended toward increased expression in *Fgf20*^*-/-*^ cochleae, albeit not significantly (padj = 0.11 and 0.38, respectively). *Bmp4* is interesting because of its importance in patterning the outer sulcus, prosensory domain, and Kölliker’s organ (Ohyama et al. 2010).

Examining the expression of these genes by ISH did not reveal noticeable changes between *Fgf20*^*-/+*^ and *Fgf20*^*-/-*^ cochleae. We hypothesize that because TRAP depletes for the outer sulcus and roof of the cochlea, TRAPseq is highly sensitive to the expression of markers of these regions in the prosensory domain. Therefore, TRAPseq may be much more sensitive than ISH to detect small changes in the expression of genes such as *Gata2, Meis2*, and *Lmx1a*, which may represent a shift in the boundary between the prosensory domain and outer sulcus.

### Cell cycle regulators are downregulated in *Fgf20*^*-/-*^ cochleae

TRAPseq revealed many differentially expressed cell cycle regulators. We have shown before that *Fgf20* by itself does not appear to regulate the cell cycle or proliferation in the developing cochlea. However, *Fgf20* is redundant with *Fgf9* in indirectly regulating prosensory progenitor proliferation at earlier developmental stages (E11.5-E12.5) (Huh et al. 2015) and is redundant with *Sox2* in regulating Kölliker’s organ proliferation at E14.5 (Yang et al. 2019). Therefore, we believe the finding of differentially expressed cell cycle regulators may be reflective of the redundant and stage-specific functions of *Fgf20* in regulating proliferation. As expected, while *Cdc20* conditional-null cochleae are short, they do not exhibit the HC differentiation or patterning defect found in *Fgf20*^*-/-*^ cochleae. We conclude that the ∼10-20% decrease in length of *Fgf20*^*-/-*^ cochleae may be attributable to decreased expression of these cell cycle regulators in sensory progenitors. It is also possible that *Fgf20* has a previously unidentified role in regulating prosensory cell cycle exit, and decreased expression of these cell cycle regulator genes are reflective of premature cell cycle exit.

### *Fgf20* regulates *Sall1*, a gene implicated in human sensorineural hearing loss

We found that members of the *Sall* family, *Sall1, Sall2*, and *Sall3* are expressed in the prosensory domain at E14.5. *Sall1, Sall3*, and potentially *Sall2* showed decreased expression by TRAPseq and ISH in *Fgf20*^*-/-*^ cochleae, suggesting that they may be regulated by FGF20 signaling. Notably, in the kidney, *Sall1* expression has been shown to be regulated by FGF signaling (Poladia et al. 2006). As mentioned previously, mutations in *SALL1* causes Townes-Brocks syndrome (TBS) in humans, an autosomal dominant disorder with variable presentation of phenotypes including sensorineural hearing loss (Kohlhase et al. 1998). *Sall1*^*Δ/+*^ mice mimic TBS, including hearing loss (Kiefer et al. 2003). However, whether cochlea development is affected in these mice has not been studied. We decided to examine *Sall1*^*Δ*^ cochleae due to evidence suggesting that the truncated Sall1^Δ^ protein acts as a dominant negative on other members of the Sall family (Kiefer et al. 2003, 2008).

We found that *Sall1*^*Δ/+*^ had normal HC patterning, but exhibited a small decrease in the total number of OHCs. This is reminiscent of the *Fgf20*^*-/-*^ and *Fgf20*;*Sox2* compound mutant phenotypes, in which OHCs are the most sensitive to the loss of FGF20 (Huh et al. 2012; Yang et al. 2019). We are not sure, however, how much this reduction in the number of OHCs contributes to the hearing defect found in these mice.

*Sall1*^Δ/Δ^ cochleae exhibited a more severe defect than *Sall1*^*Δ/+*^ cochleae, including shorter cochlea length, a small decrease in the total number of IHCs, and a large decrease in the total number of OHCs. We do not know whether the decrease in HC number is solely attributable to the shorter cochlea length. It is possible that both the HC and cochlea length phenotypes are the result of a defect in prosensory progenitor proliferation, such as that found in *Fgf20*;*Fgf9* double mutant mice (Huh et al. 2015), or the result of a defect in prosensory specification, such as that found in *Sox2* mutant mice (Kiernan et al. 2005). It is also possible that the decrease in HC number is due to a defect in differentiation, similar to that found in *Fgf20*^*-/-*^ cochleae.

Interestingly, *Sall1*^Δ/Δ^ cochleae also appeared to exhibit a delay in the apical progression of HC maturation, similar to *Fgf20*^-/-^ cochleae (Huh et al. 2012). Furthermore, *Sall1*^Δ/Δ^ cochleae contained numerous ectopic IHCs, found outside of the normal row of IHCs, a patterning defect that again is reminiscent of *Fgf20*^*-/-*^ and *Fgf20*;*Sox2* compound mutant phenotypes (Huh et al. 2012; Yang et al. 2019). The interaction between *Fgf20, Sox2*, and *Sall1/3* is a topic to explore in future studies. Based on all of these results, we conclude that the decreased expression of *Sall1* and *Sall3* may contribute to the OHC and patterning defects found in *Fgf20*^*-/-*^ cochleae.

## Conclusions

The *Fgf20*^*-/-*^ cochlea phenotype includes loss of two-thirds of OHCs, abnormal patterning of the remaining HCs, shorter cochlea length, premature onset of differentiation, and delayed apical progression of differentiation and maturation. Here, we did not identify one single gene that can account for the majority of the *Fgf20*^*-/-*^ phenotype. However, we identified many FGF20-regulated genes that may contribute to parts of the phenotype. For instance, *Hey1, Hey2*, and possibly *Heyl* may account for the premature onset of differentiation phenotype; *Sall1, Sall3*, and *Fat3* may account for the OHC differentiation, patterning, and delay in maturation phenotypes; and cell cycle regulators such as *Cdc20* may account for the progenitor proliferation phenotypes in *Fgf20*;*Fgf9* and *Fgf20*;*Sox2* compound mutants. We conclude that the dramatic *Fgf20*^*-/-*^ phenotype in which gaps in the sensory epithelium separate islands of HCs and SCs may not be explained by a straightforward lateral compartment differentiation defect. Rather, the phenotype may be the result of disruptions to a combination of FGF20-regulated processes, including prosensory progenitor proliferation, differentiation, maturation, and timing of differentiation. Given the complexity of organ of Corti development, we hypothesize that small disturbances to such processes can lead to much larger defects in overall development.

## EXPERIMENTAL PROCEDURES

### Mice

All studies performed were in accordance with the Institutional Animal Care and Use Committee at Washington University in St. Louis (protocol #20190110 and #20170258).

Mice were group housed with littermates, in breeding pairs, or in a breeding harem (2 females to 1 male), with food and water provided ad libitum. Mice were of mixed sexes and maintained on a mixed C57BL/6J x 129×1/SvJ genetic background, except *Sall1*^*Δ*^ mice, which were maintained on an ICR genetic background. The following mouse lines were used:

- *Fgf20*^*Cre*^: knockin allele containing a sequence encoding a GFP-Cre fusion protein replacing exon 1 of *Fgf20*, resulting in a null mutation (Huh et al. 2015).
- *Fgf20*^*βgal*^: knockin allele containing a sequence encoding β-galactosidase (βgal) replacing exon 1 of *Fgf20*, resulting in a null mutation (Huh et al. 2012).
- *ROSA*^*fsTRAP*^: knockin allele containing a loxP-Stop-loxP sequence followed by a sequence encoding L10a-eGFP, targeted to the ubiquitously expressed ROSA26 locus. Upon Cre-mediated recombination, the polysomal protein L10a-eGFP is expressed (Zhou et al. 2013).
- *ROSA*^*mTmG*^: knockin allele containing a sequence encoding a membrane-localized tdTomato (mT) flanked by loxP sequences, followed by a sequence encoding a membrane-localized eGFP (mG), targeted to the ubiquitously expressed *ROSA26* locus. In the absence of Cre-mediated recombination, mT is expressed; upon Cre-mediated recombination, mG is alternatively expressed (Muzumdar et al. 2007).
- *Sall1-ΔZn*^*2-10*^ (*Sall1*^*Δ*^): mutant allele expressing a truncated Sall1 protein designed to mimic a mutation that causes Townes-Brocks syndrome (Kiefer et al. 2003).
- *Cdc20*^*flox*^: allele containing loxP sequences flanking exon 2 of *Cdc20*; upon Cre-mediated recombination, results in a null allele (Manchado et al. 2010).

### Translating ribosome affinity purification (TRAP)

Affinity matrix preparation: for each immunoprecipitation (IP): 30 µl of Streptavidin MyOne T1 Dynabeads (Invitrogen, 65602) were washed in 1x PBS using an end-over-end tube rotator and a magnet, and resuspended in 88 µl of 1x PBS and conjugated to 12 µl of 1 µg/µl biotinylated protein L (Pierce 29997) in PBS for 35 min at room temperature (RT) with gentle end-over-end mixing on a tube rotator. Conjugated beads were then washed with 1x PBS + 3% IgG and protease-free BSA (Jackson ImmunoResearch, 001-000-162) x5, followed by three washes in low-salt buffer (20 mM HEPES KOH, pH 7.4, 10 mM MgCl_2_, 150 mM KCl, 1% NP-40 [Sigma I8896-50ML], 0.5 mM DTT [Sigma, 646563], 100 µg/ml cycloheximide [Sigma C4859-1ML]). Conjugated beads were then resuspended in low-salt buffer and mixed with 50 µg each of anti-GFP antibodies Htz-GFP-19C8 and Htz-GFP-19F7 (Memorial Sloan-Kettering Monoclonal Antibody Facility) overnight at 4°C with gentle end-over-end mixing to make the affinity matrix. Immediately before IP, the affinity matrix was washed in low-salt buffer x3.

Sample collection: E14.5 embryos were harvested, on ice, from a mating producing a 1:1 ratio of *Fgf20*^*Cre/+*^;*ROSA*^*fsTRAP/+*^ and *Fgf20*^*Cre/βgal*^;*ROSA*^*fsTRAP/+*^ progeny. Embryos were staged based on vaginal plug (E0.5 at noon on the day plug is found) and on interdigital webbing. Embryos with too much or too little interdigital webbing were not harvested. Embryos were genotyped by LacZ staining to look for *Fgf20*^*βgal*^ expression in back skin hair follicles (back skin from embryos were incubated in 2 mM MgCl_2_, 5 mM K3, 5 mM K4, 0.02% NP-40, and 1 mg/ml X-gal in N,N-dimethylformamide in 1x PBS for 30 min at 37°C, protected from light). Ventral otocysts from the embryos were dissected out in dissection buffer (1x HBSS, 2.5 mM HEPES-KOH, pH 7.4, 35 mM glucose, 4 mM NaHCO_3_, 100 µg/ml cycloheximide), separated from the dorsal otocyst (vestibule) without removal of the otic capsule, and pooled together by genotype. Each sample contained pooled ventral otocysts from 3-7 embryos. Pooled ventral otocysts were homogenized in lysis buffer (20 mM HEPES KOH, pH 7.4, 150 mM KCl, 10 mM MgCl_2_, EDTA-free protease inhibitors [Roche, 04693159001], 0.5 mM DTT, 100 µg/ml cycloheximide, 10 µl/ml rRNasin [Promega N2515], 10 µl/ml Superasin [Applied Biosystems, AM2696]) using a pre-chilled Kontes homogenizer (Kontes, 885512-0020). To remove the nuclear fraction, homogenized samples were centrifuged for 10 min at 2000 g, 4°C. The supernatant (S2) was mixed with 1/8 volume of 10% NP-40 and 300 mM DHPC (reconstituted in lysis buffer; Avanti Polar Lipids 850306P) and incubated for 10 min on ice. To remove the mitochondrial fraction, samples were then centrifuged for 15 min at 20,000 g, 4°C. 60 µl of the supernatant (S20) was saved as the pre-IP (pre-TRAP) control. The pre-TRAP S20 samples were incubated at 4°C until the RNA purification step, which was done in conjunction with TRAP samples. The rest of the S20 was used for IP.

Immunoprecipitation: S20 was mixed with the affinity matrix for 24 hours at 4°C with end-over-end mixing. The mixture (TRAP sample) was washed in high-salt buffer (20 mM HEPES KOH, pH 7.4, 10 mM MgCl_2_, 350 mM KCl, 1% NP-40, 0.5 mM DTT, 100 µg/ml cycloheximide, 1 µl/ml rRNasin, 1 µl/ml Superasin) for 2 min at RT, x4.

RNA purification: the Arcturus Picopure RNA Isolation Kit (Thermo Fisher, 12204-01) was used to isolate RNA from pre-TRAP and TRAP samples according to manufacturer’s instructions. RNA was eluted in 13 µl of elution buffer. Ventral otocysts from 3-7 embryos ranged between 4-20 ng of TRAP RNA. RNA samples were stored at −80°C until use in downstream applications.

### Quantitative RT-PCR

cDNA was synthesized from pre-TRAP and TRAP RNA using the iScript Select cDNA Synthesis Kit (Bio-Rad, #170-8841). mRNA expression was measured using TaqMan Fast Advanced Master Mix (Life Technologies, 4444557) and TaqMan assay probes for *Twist2* and *Id2. Gapdh* was used as normalization control. Results were analyzed by the ΔΔCT method (normalized to *Gapdh*, then normalized to pre-TRAP). Each sample represents TRAP RNA from one litter.

### cDNA library preparation and sequencing

cDNA library preparation and sequencing were done at the Genome Technology Access Center (GTAC) at Washington University (gtac.wustl.edu). RNA samples were analyzed on an Agilent 2100 Bioanalyzer; all sequenced RNA samples had an RNA Integrity Number (RIN) of ≥ 8.8. Clontech SMARTer kit was used for cDNA library preparation and amplification. The TRAPseq results presented are from two sequencing experiments. cDNA library preparation was done independently in the two experiments. In both experiments, 8 TRAP samples (4 *Fgf20*^*-/+*^ and 4 *Fgf20*^*-/-*^) and 4 pre-TRAP samples (2 *Fgf20*^*-/+*^ and 2 *Fgf20*^*-/-*^) were sequenced on one Illumina HiSeq 3000 lane, with single reads, 1 × 50 bps. 24 samples were sequenced in total between the two experiments (12 samples multiplexed per lane per experiment). Sequencing produced between 22 and 38 million reads per sample.

### Bioinformatic analysis

#### Basecalling was performed with Illumina RealTimeAnalysis software

The resulting bcl files were demultiplexed with Illumina’s bclToFastq2. Both steps were performed by GTAC.

#### Alignment

Reads were mapped to GRCm38.p5 (Ensemble, GCA_000001635.7) (Howe et al. 2020) using STAR (Dobin et al. 2013), with the GRCm38.91 annotation file (Ensembl). Default parameters were used, except for the following: multi-sample 2-pass, with default settings on first pass and sjdbFileChrStartEnd (for novel splice junctions), ScoreMinOverLread=0.4, MatchNminOverLread=0.4, MismatchNmax=5 on second pass (these parameters gave the most consistent unmapped reads % across all 24 TRAPseq samples). 95-99.5% of reads were mapped per sample.

#### Counting and DEG analysis

Analyses were performed in R using packages from Bioconductor (bioconductor.org). BAM files were indexed and sorted using Rsamtools (Morgan et al. 2018). Gene models were defined using the GRCm38.91 annotation file (Ensembl) with GenomicFeatures (Lawrence et al. 2013). Reads were counted using the SummarizeOverlaps method (mode = Union) from the package GenomicAlignments (Lawrence et al. 2013). Genes were filtered out from downstream analysis if less than 8 of 24 samples had 25 or more reads. PC analysis showed separation between the 8 pre-TRAP samples and 16 TRAP samples along PC1, and also separation between sequencing experiment 1 and experiment 2 along PC2. Removal of Unwanted Variation from RNA-Seq Data (RUVSeq) (Risso et al., 2014) was used to correct for this batch effect (RUVs function, k = 1). DESeq2 (Love et al. 2014, 2) with RUVs correction factors was used for DEG analysis, with alpha = 0.1 and Benjamini-Hochberg multiple-comparisons correction.

#### Pathway analysis

gene ontology (GO) analysis was done using the Bioconductor package topGO (Alexa and Rahnenfuhrer 2016) with the following parameters: nodeSize = 10; ontology 30 = biological processes (BP); algorithm = elim; statistic = fisher’s exact test. Protein-protein interaction network analysis was performed using STRING version 11.0 (Snel et al. 2000; Szklarczyk et al. 2019) with the following parameters: active interaction sources include textmining, experiments, databases, co-expression, neighborhood, gene fusion, co-occurrence; minimum required interaction score = high confidence (0.700).

### Sample preparation and sectioning for histology and in situ hybridization

For whole mount cochleae, inner ears were dissected out of P0 pups and fixed in 4% PFA in PBS overnight at 4°C with gentle agitation. Samples were then washed x3 in PBS. Cochleae were dissected away from the vestibule, otic capsule, and periotic mesenchyme with Dumont #55 Forceps (RS-5010, Roboz, Gaithersburg, MD). The roof of the cochlear duct was opened up by dissecting away the stria vascularis and Reissner’s membrane; tectorial membrane was removed to expose hair and supporting cells.

For sectioning, heads from E14.5 embryos were fixed in 4% PFA in PBS overnight at 4°C with gentle agitation. Samples were then washed x3 in PBS and cryoprotected in 15% sucrose in PBS overnight and then in 30% sucrose in PBS overnight. Samples were embedded in Tissue-Tek O.C.T. compound (4583, VWR International, Radnor, PA) and frozen on dry ice. Serial horizontal sections through base of the head were cut at 12 µm with a cryostat, dried at room temperature, and stored at −80°C until use.

### RNA in situ hybridization

Probe preparation: mouse cDNA plasmids containing the following inserts were used to make RNA in situ probes, and were cut and transcribed with the indicated restriction enzyme (New England Biolabs) and RNA polymerase (New England Biolabs): *Dusp6* (412 bp, Acc65I, T7, gift of Suzanne Mansour), *Etv1* (2500 bp, SpeI, T7, gift of Sung-Ho Huh), *Spry1* (1500 bp, EcoRI, T7, gift of George Minowada), *Spry4* (900 bp, EcoRI, T7, gift of George Minowada), *Tectb* (2746 bp, EcoRI, T7, gift of Doris Wu), *Tecta* (4382 bp, NotI, T7, gift of Doris Wu), *Epyc* (1522 bp, EcoRI, T7, Image clone 4037028), *Fat3* (945 bp, EcoRI, T7, gift of Lisa Goodrich), *Heyl* (1895, BamHI, T7, Image clone 40142873) *Sall1* (450 bp, HindIII, T7), *Sall2* (431 bp, EcoRi, T7), *Sall3* (551 bp, XbaI, T3), *Gata2* (700 bp, BamHI, T3, gift of Doris Wu), *Meis2* (∼5000 bp, EcoRI, T3, gift of Yingzi Yang), *Lmx1a* (600 bp, SphI, Sp6, gift of Doris Wu), *Bmp4* (1560 bp, AccI, T7). The *Smpx* probe was made from PCR product (gift of Jinwoong Bok) and transcribed with T7.

Frozen section in situ hybridization: frozen slides were warmed for 20 min at room temperature and then 5 min at 50°C on a slide warmer. Sections were fixed in 4% PFA in PBS for 20 min at room temperature, washed x2 in PBS and treated with pre-warmed 10 µg/ml Proteinase K (03115828001, Sigma-Aldrich, St. Louis, MO) in PBS for 7 min at 37°C. Sections were then fixed in 4% PFA in PBS for 15 min at room temperature, washed x2 in PBS, acetylated in 0.25% acetic anhydrate in 0.1M Triethanolamine, pH 8.0, for 10 min, and washed again in PBS. Sections were then placed in pre-warmed hybridization buffer (50% formamide, 5x SSC buffer, 5 mM EDTA, 50 µg/ml yeast tRNA) for 3 h at 60°C in humidified chamber for prehybridization. Sections were then hybridized in 10 µg/ml probe/hybridization buffer overnight (12-16 h) at 60°C. The next day, sections were washed in 1x SSC for 10 min at 60°C, followed by 1.5x SSC for 10 min at 60°C, 2x SSC for 20 min at 37°C x2, and 0.2x SSC for 30 min at 60°C x2. Sections were then washed in KTBT (0.1 M Tris, pH 7.5, 0.15 M NaCl, 5 mM KCl, 0.1% Triton X-100) at room temperature and blocked in KTBT + 20% sheep serum + 2% Blocking Reagent (11096176001, Sigma-Aldrich, St. Louis, MO) for 4 h. Blocking Reagent was dissolved in 100 mM Maleic acid, 150 mM NaCl, pH 7.5. Sections were then incubated in sheep anti-Digoxigenin-AP, Fab fragments (1:1000, 11093274910, Sigma-Aldrich, St. Louis, MO) in KTBT 14 + 20% sheep serum + 2% Blocking Reagent overnight at 4°C. Sections were then washed x3 in KTBT for 30 min at room temperature, and then washed x2 in NTMT (0.1 M Tris, pH 9.5, 0.1 M NaCl, 50 mM MgCl_2_, 0.1% Tween 20) for 15 min. Sections were next incubated in NTMT + 1:200 NBT/BCIP Stock Solution (11681451001, Sigma-Aldrich, St. Louis, MO) in the dark at room temperature until color appeared. Sections were then washed in PBS, post-fixed in 4% PFA in PBS for 15 min and washed x2 in PBS. Finally, sections were dehydrated in 30% and then 70% methanol, 5 min each, followed by 100% methanol for 15 min. Sections were then rehydrated in 70% and 30% methanol and then PBS, 5 min each, and mounted in 95% glycerol.

### Immunofluorescence

Whole mount: cochleae were incubated in PBS + 0.5% Tween-20 (PBSTw) for 1 h to permeabilize. Cochleae were then blocked using PBSTw + 5% donkey serum for 1 h and then incubated in PBSTw + 1% donkey serum with the primary antibody overnight at 4°C. Cochleae were then washed x3 in PBS and incubated in PBS + 1% Tween-20 with the secondary antibody. After wash in PBS x3, cochleae were mounted in 95% glycerol with the sensory epithelium facing up.

Frozen slides were warmed for 30 min at room temperature and washed in PBS before incubating in PBS + 0.5% Triton X-100 (PBST) for 1 h to permeabilize the tissue. Sections were then blocked using in PBST + 5% donkey serum for 1 h and then incubated in PBST + 1% donkey serum with the primary antibody overnight at 4°C in a humidified chamber. Sections were then washed x3 in PBS and incubated in PBS + 1% Triton X-100 with the secondary antibody. After wash in PBS x3, slides were mounted in VectaShield antifade mounting medium with DAPI (H-1200, Vector Labs, Burlingame, CA).

The following compounds and antibodies were used:

- Alexa Fluor 488-conjugated Phalloidin (1:50, A12379, Invitrogen, Carlsbad, CA)
- Rabbit anti-P75NTR (1:300, AB1554, EMD Millipore, Burlington, MA)
- Alexa Fluor 555 goat anti-rabbit IgG (1:500, A21428, Invitrogen, Carlsbad, CA)

### Imaging

Light microscopy: slides were scanned using a Hamamatsu NanoZoomer slide scanning system with a 20x objective. Images were then processed with the NanoZoomer Digital Pathology (NDP.view2) software. 3D specimens were imaged using an Olympus SZXZ110 stereo microscope equipped with an Olympus DP70 camera.

Fluorescent microscopy was done using a Zeiss LSM 700 confocal or Zeiss Axio Imager Z1 with Apotome 2, with z-stack step-size determined based on objective lens type (10x or 20x), as recommended by the ZEN software (around 1 µm). Fluorescent images shown are maximum projections. Images were processed with ImageJ (imagej.nih.gov).

### Image analysis and quantification

Measurements and cell quantification (using the Cell Counter plugin by Kurt De Vos) were done using ImageJ and Fiji (Schindelin et al. 2012). Total cochlear duct length was defined as the length from the very base of the cochlea to the very tip of the apex, along the tunnel of Corti, measured on whole-mount cochlea. Hair cells and stereocilia bundles were identified via Phalloidin, which binds to F-actin (Avinash et al. 1993). Inner pillar cells were labeled via P75NTR (Mueller et al. 2002). Inner hair cells (IHCs) were differentiated from outer hair cells (OHCs) based on their neural/abneural location, respectively, relative to P75NTR-expressing inner pillar cells. For total cell counts, IHCs and OHCs were counted along the entire length of the cochlea.

### Statistical analysis and plotting

All figures were made in Canvas X (ACD systems). RNA sequencing data analysis and plotting, were performed using R (r-project.org) in R studio (rstudio.com) PCA graphs were made using the plotPCA function from the package RUVSeq; volcano plots were made using modified code from Stephen Turner (gist.github.com/stephenturner). See Bioinformatic analysis section for more details on RNA sequencing data analysis. All other data analysis and plotting were performed using Python (python.org) in Jupyter Notebook (jupyter.org) with the following libraries: Pandas (pandas.pydata.org), Seaborn (seaborn.pydata.org), NumPy (numpy.org) and SciPy (scipy.org). Plotting was done using the Matplotlib library (matplotlib.org). Statistics (t-test, one-way ANOVA, and two-way ANOVA) were performed using the SciPy module Stats; Tukey’s HSD was performed using the Statsmodels package (statsmodels.org). All comparisons of two means were performed using two-tailed, unpaired Student’s t-test. For comparisons of more than two means, one-way ANOVA was used. For significant ANOVA results at α = 0.05, Tukey’s HSD was performed for post-hoc pair-wise analysis. In all cases, p < 0.05 was considered statistically significant. All statistical details can be found in the figures and figure legends. In all cases, each sample (each data point in graphs) represents one animal. Based on similar previous studies, a sample size of 3-5 was determined to be appropriate. Error bars represent mean ± standard deviation. For qualitative comparisons (comparing expression via immunofluorescence or RNA in situ hybridization), at least three samples were examined per genotype. All images shown are representative. No data were excluded from analysis.

## ACKNOWLEDGEMENTS

We thank J. Dougherty and his laboratory for their help with the TRAP technique, and B. Zhang (Department of Developmental Biology), and T. Sinnwell and M. Heinz (Genome Technology Access Center in the Department of Genetics) for their help with RNA sequencing and analysis. We also thank Cole Ferguson for supplying us with *Cdc20*^*flox*^ mice. This work was funded by the Department of Developmental Biology at Washington University, NIH/National Institute on Deafness and Other Communication Disorders grant DC017042 (DMO), the Washington University Institute of Clinical and Translational Sciences which is, in part, supported by the NIH/National Center for Advancing Translational Sciences, CTSA grant UL1TR002345 (JIT471 to DMO), March of Dimes grant 6-FY13-127 (MR), and the Rare Disease Foundation/BC Children’s Hospital Foundation. GTAC is partially supported by NCI Cancer Center Support Grant P30 CA91842 to the Siteman Cancer Center and by ICTS/CTSA Grant UL1TR000448 from the National Center for Research Resources (NCRR). HOPE Center Alafi Neuroimaging Laboratory is supported by NCRR grant 1S10RR027552, the Auditory and Vestibular Microscopy and Digital Imaging Core is supported by NIH grant P30 DC004665. The authors have no conflicts of interest to disclose.

## AUTHOR CONTRIBUTIONS

Conceptualization, L.M.Y., M.R., and D.M.O.; Methodology, L.M.Y. and D.M.O.; Formal Analysis, L.M.Y.; Investigation: L.M.Y. and L.S.; Resources: M.R. and D.M.O.; Writing – Original Draft: L.M.Y; Writing – Review & Editing: L.M.Y., M.R., and D.M.O.; Supervision: D.M.O.; Funding Acquisition: M.R. and D.M.O.

## Data availability

The data discussed in this publication have been deposited in NCBI’s Gene Expression Omnibus (Edgar et al. 2002) and are accessible through GEO Series accession number GSE148380 (https://www.ncbi.nlm.nih.gov/geo/query/acc.cgi?acc=GSE148380).

## Supplemental files

S1 – preTRAP vs. TRAP DEG analysis

S2 – *Fgf20*^*-/+*^ vs. *Fgf20*^*-/-*^ DEG analysis

## List of abbreviations

DEG: differentially expressed gene
GER: greater epithelial ridge
HC: hair cell
IHC: inner hair cell
IP: immunoprecipitation
KO: Kölliker’s organ
LER: lesser epithelial ridge
OC: organ of Corti
OHC: outer hair cell
OS: outer sulcus
padj: adjusted p-value
PCA: principal component analysis
PD: prosensory domain
SC: supporting cell
TBS: Townes-Brocks syndrome
TRAP: translating ribosome affinity purification
TRAPseq: TRAP combined with next generation mRNA sequencing

